# Developing a Multiscale Neural Connectivity Knowledgebase of the Autonomic Nervous System

**DOI:** 10.1101/2024.10.25.620360

**Authors:** Fahim T. Imam, Thomas H. Gillespie, Ilias Ziogas, Monique C. Surles-Zeigler, Susan Tappan, Burak I. Ozyurt, Jyl Boline, Bernard de Bono, Jeffrey S. Grethe, Maryann E. Martone

## Abstract

The Stimulating Peripheral Activity to Relieve Conditions (SPARC) program is a U.S. National Institutes of Health (NIH) funded effort to enhance our understanding of the neural circuitry responsible for visceral control. SPARC’s mission is to identify, extract, and compile our overall existing knowledge and understanding of the autonomic nervous system (ANS) connectivity between the central nervous system and end organs. A major goal of SPARC is to use this knowledge to promote the development of the next generation of neuromodulation devices and bioelectronic medicine for nervous system diseases. As part of the SPARC program, we have been developing SCKAN, a dynamic knowledge base of ANS connectivity that contains information about the origins, terminations, and routing of ANS projections. The distillation of SPARC’s connectivity knowledge into this knowledge base involves a rigorous curation process to capture connectivity information provided by experts, published literature, textbooks, and SPARC scientific data. SCKAN is used to automatically generate anatomical and functional connectivity maps on the SPARC portal.

In this article, we present the design and functionality of SCKAN, including the detailed knowledge engineering process developed to populate the resource with high quality and accurate data. We discuss the process from both the perspective of SCKAN’s ontological representation as well as its practical applications in developing information systems. We share our techniques, strategies, tools and insights for developing a practical knowledgebase of ANS connectivity that supports continual enhancement.

## 1 INTRODUCTION

An NIH Common Fund initiative, the Stimulating Peripheral Activity to Relieve Conditions (SPARC, https://commonfund.nih.gov/sparc), is a large-scale program aiming to accelerate the development of therapeutic devices that modulate electrical activity in nerves to improve organ function. SPARC aims to foster the creation of robust scientific and technological foundations, paving the way for the development of more effective and targeted bio-electronic neuromodulation therapies for a wide range of diseases and conditions. The overarching goal of the first phase of the SPARC program (2017-2022) was to deliver detailed, integrated multi-species functional and anatomical maps and models of the autonomic nervous system (ANS) and its innervation of end organs. The second phase of SPARC (2022-2025) will deliver detailed anatomical and functional maps of the human vagus, as well as open-source libraries for new neuromodulation devices.

The SPARC Data and Resource Center (DRC) develops the infrastructure and standards for storing, reusing, and integrating data and other publicly available resources across the consortium. The DRC supports the data sharing platform, standards, the SPARC knowledge graph, connectivity maps, and simulation tools for the mammalian nervous system (Osanlouy et al., 2021; Surles-Zeigler et al., 2022). The SPARC Portal (https://sparc.science/) is the main entry point to SPARC data, maps, tools and community. The portal supports multiple search options across data, models, and simulations including faceted search, keyword search, and map-based query.

A key component of the SPARC project, the SPARC Connectivity Knowledgebase of the Autonomic Nervous System (SCKAN, RRID:SCR 026088), is a semantic store containing detailed, formal knowledge about ANS connectivity between the central nervous system and end organs. SCKAN serves as a powerful resource to populate, discover, and query ANS connectivity over multiple scales, species, and organs. SCKAN is being used within SPARC to generate interactive connectivity maps available via the SPARC Portal (https://sparc.science/apps/maps?id=7d8ad680). Figure 1 provides an example of SPARC’s anatomical connectivity map showing different connectivity pathways in rats sourced from SCKAN.

**Figure 1.**
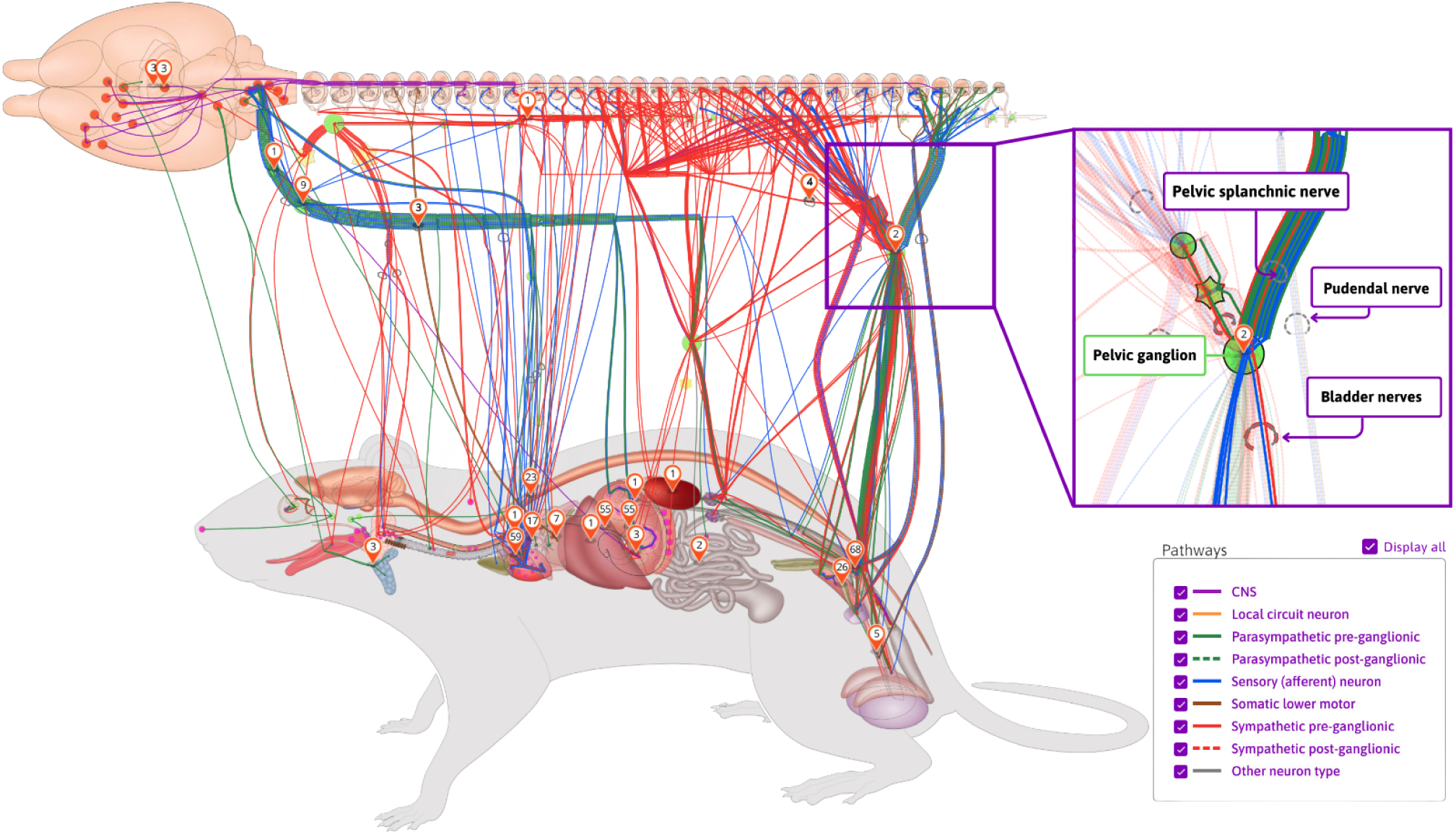
An example of SPARC’s anatomical connectivity map. These dynamic maps can be accessed via the SPARC Portal (https://sparc.science/apps/maps?id=7d8ad680). The connections pictured here are drawn directly from the SCKAN knowledgebase. The maps show the origins, terminations, and the nerves through which they travel (inset on right). Nerves are color-coded according to their phenotypes and circuit roles.

In this article, we provide an overview of SCKAN and present details about its organization, curation, and content population. We then demonstrate how SCKAN can be used to answer questions about ANS connectivity, including detailed routing of axons within individual nerves.

## 2 MATERIALS AND METHODS

We first give an overview of how connectivity information is modeled within SCKAN and then go through the steps of identifying and encoding connections to populate SCKAN. Finally, we provide details on how to access SCKAN.

### 2.1 Modeling Connectivity

SCKAN models connectivity at the neuronal level, where connections are represented based on the locations of neuronal cell bodies, axon segments, dendrites, and axon terminals. Each connection represents a population of PNS neurons with a given PNS phenotype (e.g. sympathetic pre-ganglionic) that projects to a given set of targets, where the exact cell type identity may not be known. By modeling neuron populations in terms of their connected regions, SCKAN enables the assignment of phenotypes, such as anatomical locations, to individual parts of neurons. In SCKAN, the connectivity of a neuron population is therefore modeled as ‘Neuron population *N* at *A* projects to *B* via *C*’, where *A* and *B* denote sets of anatomical regions containing the somata and axon terminals, respectively, and *C* represents the set of distinct anatomical region(s) between *A* and *B* that contain the axon segments of *N*, including nerves. Figure 2 illustrates this connectivity model for a generic neuron population. This generalized model is extended to accommodate multiple parallel locations (*A_i … l_* to *B_i … m_* via *C_i … n_*), enabling the representation of connectivity pathways that link multiple origins to multiple terminals. We use the axon sensory terminal to indicate the peripheral process of pseudo-monopolar dorsal root ganglia connections. In addition to specifying the anatomical locations of neuronal segments, each neuron population in SCKAN can be further specified with other relevant phenotypic properties, e.g.:

*•* ANS subdivision: sympathetic or parasympathetic, preganglinic or post-ganglionic
*•* Circuit role: intrinsic, motor, sensory, or projection
*•* Functional circuit role: whether the connection is excitatory or inhibitory
*•* Species and sex: the specificity of species and sex the connection was observed in

**Figure 2.**
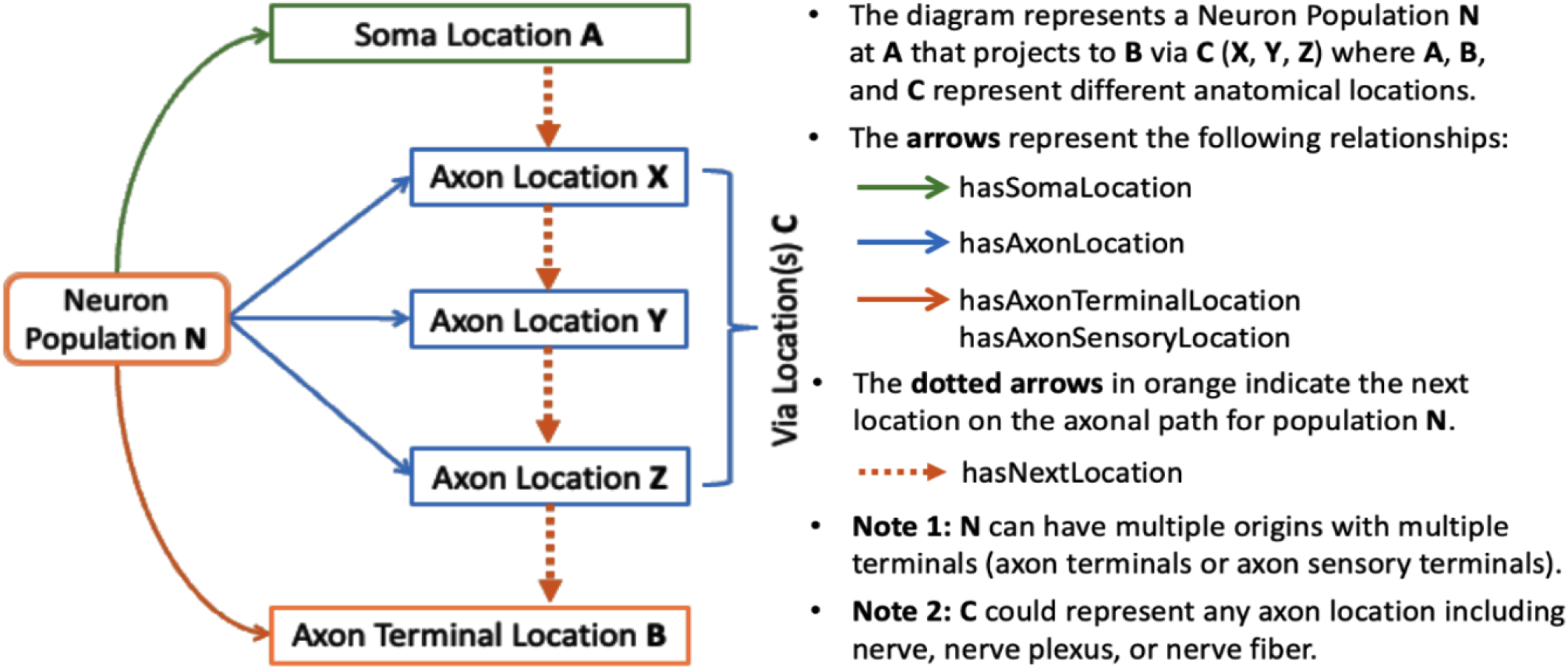
SCKAN’s connectivity model of a neuron population based on its axonal projection path.

A given population subserves a single PNS phenotype, e.g., sympathetic pre or post ganglionics, but may have multiple origins, primarily to indicate that a connection arises from multiple spinal cord segments, as well as multiple terminal locations. SCKAN currently does not model whether a neuron population projecting to multiple regions does so via branching axons or whether distinct subpopulations project to distinct targets. The connectivity of a single neuron population includes distinct anatomical structures along its projection paths from its origin(s) to its destination(s). The diagram on the left in Figure 3 shows a connectivity pathway of a parasympathetic preganglionic neuron population in SCKAN, originating at the superior salivatory nucleus and terminating at the pterygopalatine ganglion. In this example, the axon reach its target via the facial nerve, followed by the greater petrosal nerve and then the vidian nerve.

**Figure 3.**
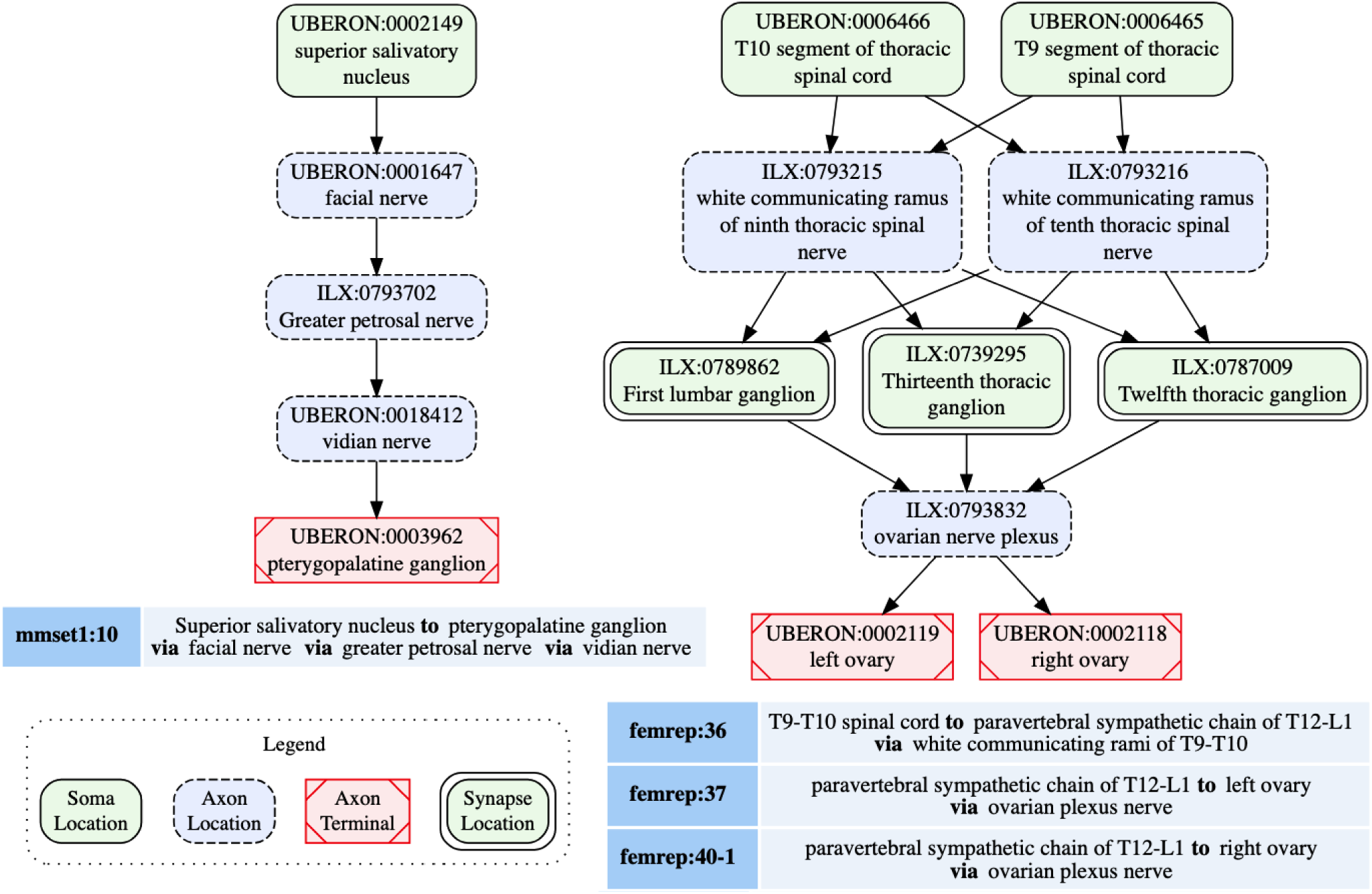
Neural connectivity examples from SCKAN. The left diagram illustrates the connectivity pathway of a parasympathetic preganglionic population, while the diagram on the right depicts a sympathetic innervation circuit of the ovary, formed by synaptic connections between pre- and postganglionic populations. Each node in the diagrams is labeled with the corresponding anatomical region, drawn from standard vocabulary sources such as UBERON or InterLex (ILX), along with its CURIE (compact URI).

SCKAN facilitates the modeling of connectivity circuits by specifying synaptic forward connections between first-order and second-order neuron populations. Preganglionic neuron populations, originating in the CNS and terminating in ANS ganglia, represent the first-order connections in SCKAN. Postganglionic populations, originating in the ANS ganglia and terminating in a specific part of a target organ, represent the second-order connections. When a forward connection is specified between two populations, the synapse location within the circuit can be identified as the common structure(s) where the preganglionic connection terminates (the location of the axon presynaptic element) and the postganglionic connection originates. The diagram on the right in Figure 3 shows an example of a sympathetic innervation circuit of the ovary, formed by the synaptic connections between a sympathetic preganglionic population (femrep:36) and two postganglionic populations (femrep:37 and femrep:40-1), observed in female rats. The preganglionic population (femrep:36) originates from two adjacent locations (T9-T10 segments of thoracic spinal cord) and terminates in three locations (T12-L1 paravertebral sympathetic chain) via the white communicating rami of T9-10. The origins (the somata locations) of the postganglionic populations (femrep:37 and femrep:40-1) are synaptically co-located with the preganglionic terminals, and their axons travel via the ovarian nerve plexus, with one terminating in the left ovary and the other in the right ovary. All the nodes representing the anatomical regions in Figure 3 are drawn from standard vocabulary sources including UBERON (RRID:SCR 010668), InterLex (ILX, RRID:SCR 016178). We discuss more about SCKAN’s vocabulary management in Section 2.4.2.

#### Extracting Connectivity Knowledge

Connectivity knowledge contained in SCKAN is derived from two principal sources: (a) Expert-contributed connectivity models, and (b) Literature- or textbook- extracted connectivity. Figure 4 provides a high-level depiction of SCKAN’s knowledge construction flow, starting with the extraction of connectivity knowledge from these sources, followed by the curation and transformation of this knowledge into formal representations, and finally, making the knowledgebase available via different graph database endpoints. The following sections detail the steps involved in this knowledge construction process.

**Figure 4.**
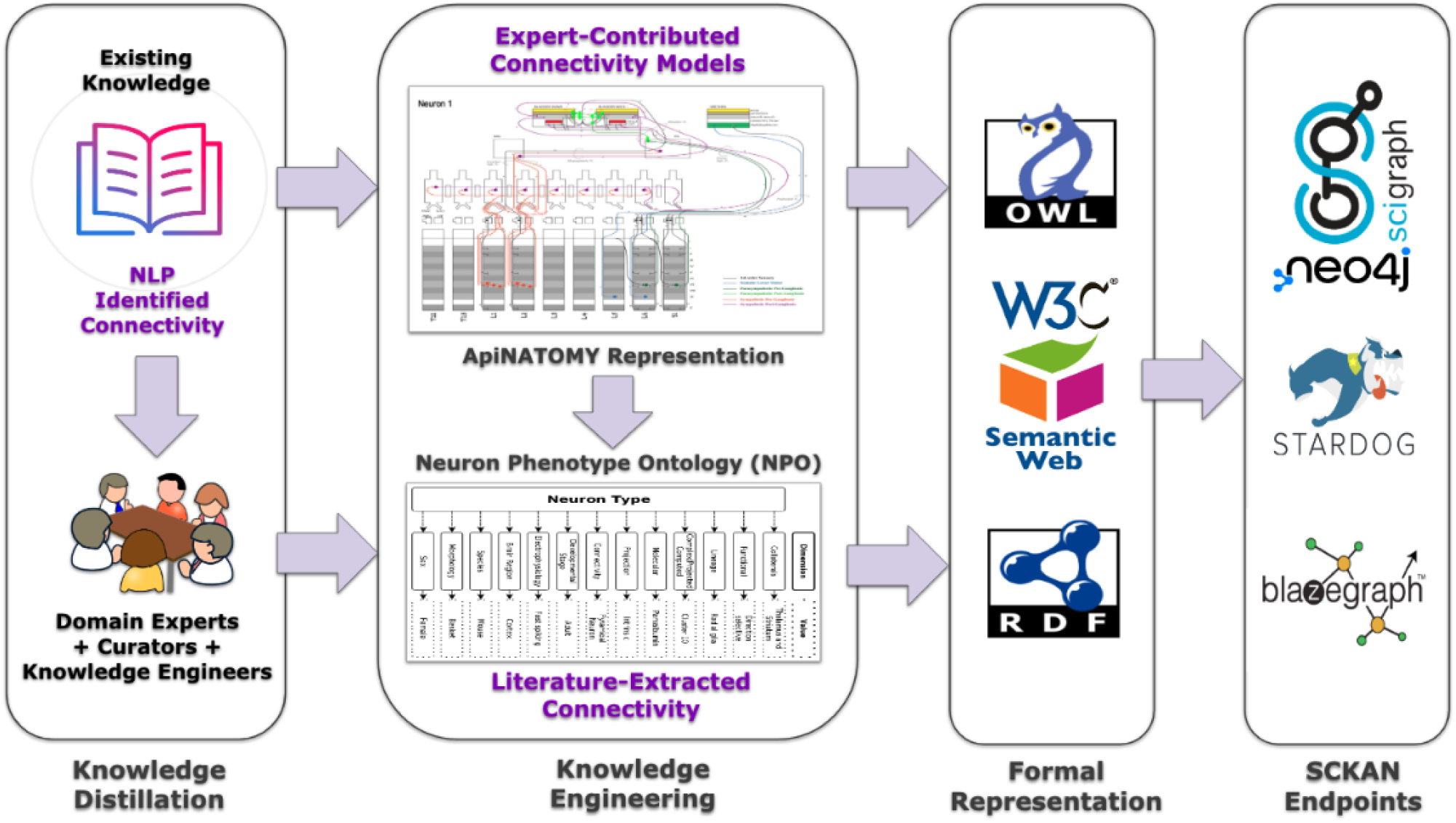
A high-level depiction of SCKAN’s knowledge construction flow. Each step of the flow is described further in the text.

### 2.2 Knowledge Distillation

The initial population of SCKAN was largely driven by detailed circuitry models of autonomic nervous system (ANS) connectivity associated with specific organs, as listed in Table 1. Each model was commissioned by the SPARC project based on its relevance to SPARC Phase I and was constructed through consultations with domain experts. The SPARC Anatomical Working Group (SAWG, RRID:SCR 026087, https://scicrunch.org/sawg), comprising anatomical experts, provided community-supported leadership and arbitration to ensure the integrity of anatomical knowledge represented in SCKAN. The expert- contributed models listed in Table 1 were developed using ApiNATOMY (RRID:SCR 018998, https://scicrunch.org/sawg/about/ApiNATOMY), a sophisticated knowledge modeling framework designed to represent and generate anatomical connectivity and pathways as diagrams, particularly in complex biological systems (Kokash and de Bono, 2021; de Bono et al., 2022). Note that as the focus of SPARC is on peripheral connectivity of the ANS, these models generally include the amount of CNS connectivity necessary to represent interactions with ANS populations, but do not aim to be comprehensive for CNS circuitry in the spinal cord, brainstem, or beyond the higher-order structures.

**Table 1.** The list of expert-contributed ANS connectivity models in SCKAN. These models were developed using ApINATOMY representations (https://scicrunch.org/sawg/about/ApiNATOMY) described in text. The SPARC Anatomical Working Group (SAWG) provided guidance to ensure the accuracy and integrity of the anatomical knowledge represented in SCKAN.

| Model Name | Description |
| --- | --- |
| Keast Model of Bladder Innervation | Neural circuits in the central and peripheral nervous system innervating the urinary bladder and urethra. Observed predominantly in rodents. |
| SAWG Model of Bronchomotor Control | Neural circuits for visceral afferents from, and para/sympathetic effectors to, the lower airways. |
| Bolser-Lewis Model of Defensive Breathing | Neural circuits controlling defensive breathing with components under the influence of the Superior Cervical Ganglion. Observed predominantly in rodents. |
| SAWG Model of the Descending Colon | Neural circuits innervating the descending colon. Includes sympathetic and visceral sensing of a prototypical segment of distal colonic muscle layers, blood supply, Peyer's patches and enteroendocrine cells. Observed predominantly in felines. |
| UCLA Model of the Heart | Neural circuits innervating the heart observed in dogs, domestic pigs, and humans. Includes the sympathetic, parasympathetic, and intrinsic sensing of the four cardiac chambers, aorta, and pulmonary trunk. |
| SAWG Model of the Pancreas | Autonomic projections to the endocrine and exocrine components of the pancreas from circuits originating from the spinal cord (sympathetic outflow via the coeliac/superior mesenteric ganglia), brainstem (vagal outflow) and gastroduodenal myenteric plexus (trans-visceral cross-talk). |
| SAWG Model of the Stomach | Autonomic and interoceptive sensory circuits between the insula and the gastric mucosa as conveyed across brain centers such as nucleus tractus solitarius and the dorsal motor nucleus of the vagus, and onwards to the stomach via the tenth cranial nerve. Also includes circuitry that cross-talks between the longitudinal and circular muscle layers, the myenteric plexus, and the layers of the mucosa for the proximal and distal stomach. |
| SAWG Model of the Spleen Innervation | Circuitry of the autonomic modulation of lymphocytic interplay between spleen and the intestinal wall. This model focuses on coeliac/superior mesenteric outflows to splenic periaarteriolar lymphoid sheets, and the vagal influence exerted on postganglionic sympathetic neurons, as well as mesenteric lymph nodes in the small intestine. |

To ensure that SCKAN and the resulting flat maps were comprehensive of major autonomic connections, we developed an NLP-based pipeline to identify papers in the scientific literature containing connectivity information (Ozyurt et al., 2021; Menke et al., 2020). The connectivity relation extraction classifier was trained to recognize sentences containing two or more peripheral anatomical structures related through connectivity, e.g., “… stimulation of sensory afferents from the trigeminal nerve results in parasympathetic vasodilation of the cerebral vasculature via interactions with the facial nerve and SPG” (White et al., 2021). The initial ANS connectivity relation extraction was conducted across the entirety of the full-length open-access subset of PubMed Central (PMC) articles as well as additional sources to which we had full-text access through publisher agreements. The data sources include the following:

*•* Full-length open-access subset of PMC: 4 million papers (downloaded in November 2020)
*•* PubMed abstracts: 31.8 million abstracts (2021 baseline)
*•* Elsevier articles from 221 journals: 290,734 papers

The initial run resulted in several thousand sentences that had to be reviewed (see next section). After the initial run, the pipelines are run monthly on new full-length PMC open-access articles. These new publications can either be articles published since the last run or full texts that became available for older articles. Generally, these monthly runs result in fewer than 50 new sentences for review.

Connectivity sentences were imported into a custom tool, OSCAP (The Open SAWG Connectivity Authoring Pipeline). Sentences and the papers from which they were extracted were reviewed by curators to determine whether there was new connectivity information that needed to be added to SCKAN. Curators were individuals with a neuroscience background, familiar with the contents of SCKAN and basic knowledge engineering. Curators did not necessarily have a deep knowledge of the ANS, but they were competent to interpret connectivity information contained within experimental papers. Curators first determined whether the information presented within the sentence was within scope, i.e., ANS mammalian connectivity. Any sentences that were determined to have been returned in error, i.e., they did not contain connectivity information, were flagged for feedback to further train the algorithm. Relevant sentences/papers were then checked against the contents of SCKAN to see if that information was already present in the database using a query interface, SCKAN Explorer, described in Section 2.5.1. If the answer was yes, curators checked to see if new information about the connection was available, e.g., verifying an existing connection in another species. If this information was not in SCKAN, the sentence was flagged and the curator would add the connection, usually by tracking down additional references.

Once the curator identified a set of likely connections, a domain expert from SAWG reviewed the evidence and provided input on whether these connections should be added to SCKAN. Experts provided input as to whether the proposed connections were well established or, for less well known connections, if the evidence provided was strong. If the evidence was less compelling, a decision was made by the curators and experts as to whether it should be included in the knowledge base. In these cases, any projections added were flagged with an alert explaining any concerns about the nature of the evidence.

We found that the most efficient way for domain experts to review proposed additions was to distill the proposed neuron population into a single sentence of the form “Structure A projects to Structure B via route C”. This sentence was followed by evidence provided in the paper or papers on which the statement was based. In this way, the curator was explicit about the proposed connections and the expert could indicate whether they agreed with the statement (or parts of the statement) based on their expert knowledge and the evidence provided. The form of the sentence also directly mapped onto the underlying data model of SCKAN (see Section 2.1), facilitating communication with the knowledge engineers. This distillation process therefore provided the critical bridge between the domain expert and the knowledge engineer by providing the information in a way that could be easily reviewed and checked for accuracy by both.

### 2.3 Knowledge Engineering

Once the curator and domain experts agreed on the neuron populations to be added, the curator prepared them for handoff to the knowledge engineers. This process involved expressing each connection in terms of anatomical locations of soma, axon segment and terminals, mapping anatomical locations to UBERON or Interlex identifiers (see Section 2.4.2), and providing phenotype, species, and sex information. All connectivity statements were accompanied by provenance indicating their source. Acceptable sources include scientific articles, textbooks or expert knowledge. Each of these sources was mapped to the appropriate identifier: DOI or PMID, ISBN, or ORCIDs respectively. Curators could also include additional notes to any population to provide more information.

During the curation process, the curator consulted with the knowledge engineers about how to represent more challenging aspects of representation, e.g., branching, unidentified nerves. Over time, a set of rules were developed that were included in a curation manual to ensure that they were applied consistently (Link to Curation Manual). The curators were then tasked with performing initial knowledge engineering steps by mapping the entities within the distilled statements into structured knowledge statements. The majority of connectivity statements distilled by the knowledge curators and currently in SCKAN were entered using a standardized Google Sheets template, set up by the knowledge engineers (Link to an example template). This template provided a semi-formal structure for curators to record connectivity statements according to SCKAN-specific phenotypic dimensions, along with detailed provenance and curation notes for each neuron population. Curators used the commenting and tagging features of Google Sheets to consult with SAWG and knowledge engineers whenever they encountered questions regarding their entries. Each connectivity statement recorded by the curators had to be approved by SAWG before being added to SCKAN. SAWG also provided feedback on the anatomical terms used in the statements. Once curators finalized their work, the curated Google Sheets were used to automatically transform the data into formal, machine-processable connectivity statements for SCKAN. Note that the process of reviewing and curating connectivity statements has since been replaced by the SCKAN Composer tool described below.

Over time, we found that the distillation and knowledge engineering processes were error prone and time consuming, e.g., all IDs for the anatomical entities had to be supplied manually. To streamline the distillation process, we developed SCKAN Composer (RRID:SCR 024441) which supports the entire distillation and curation pipeline, replacing both OSCAP and spreadsheets, and providing support for steps such as ontology ID look-ups for anatomical entities. Composer was created to reduce the human effort required to identify, validate, and construct connectivity relationships and circuits, both initially and over time. The user interface of Composer allows the curator to draft a preliminary “A to B via C statement”, add tags, notes sections, provenance hyperlinks, species and sex identification. After initial distillation, the curator crafts the path trajectory utilizing anatomical entities retrieved from a data table that includes ontological IDs and synonyms. Composer automatically generates the stereotyped ‘A to B via C’ statement and a population diagram for review by the anatomical experts based on the entities selected by the curator. The production version of SCKAN Composer (https://composer.scicrunch.io/) has recently been released and is now being used to populate SCKAN with new connections. Further details about its development and capabilities will be included in a subsequent publication.

### 2.4 Formal Representation

Once the pre-engineering steps were completed, the populations are handed off to the knowledge engineers for formal encoding. SCKAN is formally represented as an ontology using OWL-DL (Hitzler et al., 2009), a W3C standard language for developing ontologies for the semantic web. OWL enables SCKAN to be verified for consistency, infer implicit connectivity knowledge, and classify its populations through automated reasoning. SCKAN leverages the Neuron Phenotype Ontology (NPO) as its foundational data model. NPO, developed as part of the BRAIN Initiative Cell Census Network (BICCN, RRID:SCR 015820, https://www.biccn.org/) and SPARC initiatives, provides a FAIR approach for representing and classifying neuron types based on their phenotypic properties (Gillespie et al., 2022). The notion of neuron types in NPO is generalized in SCKAN to represent region-to-region connectivity at the neuron population level. The OWL representation of a neuron population in SCKAN denotes a theoretical grouping of neurons based on shared phenotypic properties without specifying individual neuron types.

SCKAN employs a subset of phenotypic properties from NPO to define its neuron populations. Figure 5 illustrates the SCKAN-specific subset of NPO’s phenotypic properties and their possible values (property ranges). Complying to the FAIR principles (Wilkinson et al., 2016), SCKAN reuses the terms associated with its phenotypes from community ontologies and vocabulary sources such as the UBERON (RRID:SCR 010668), InterLex (ILX, RRID:SCR 016178), NCBI Taxonomy (RRID:SCR 003256), and

**Figure 5.**
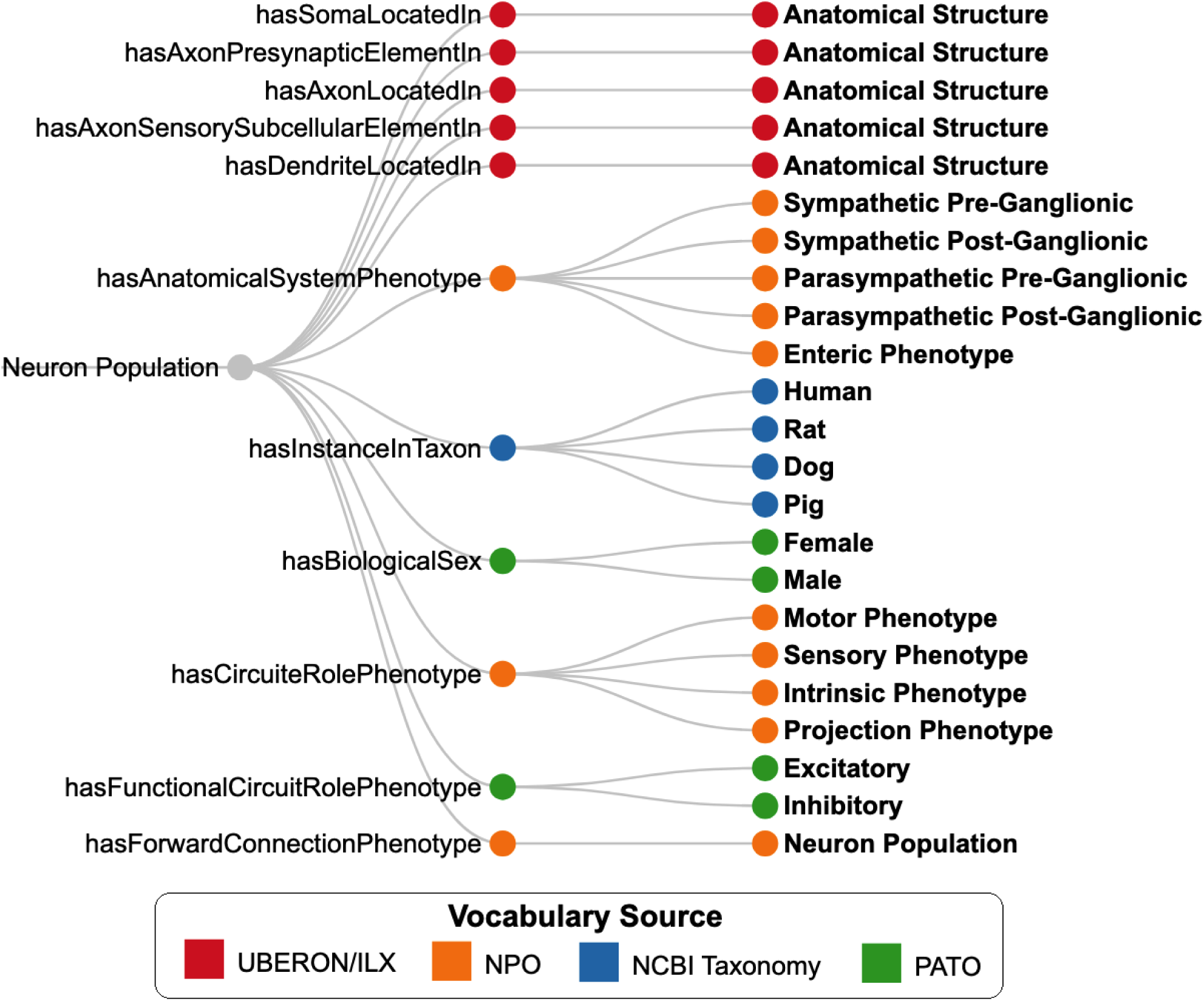
SCKAN-specific subset of properties (nodes in the middle) from the Neuron Phenotype Ontology (NPO) and their possible range of values (nodes on the right). Nodes are color-coded based on the vocabulary sources of the phenotype values.

PATO (RRID:SCR 004782), as indicated by distinct node colors in the figure. Anatomical entities required to model locational phenotypes are primarily sourced from UBERON, an integrative multi-species anatomy ontology (Mungall et al., 2012). Refer to Section 2.4.2 for more details on vocabulary management.

A class representing a neuron population in SCKAN must have its locational phenotypes associated with at least one origin and one destination. Each neuron population class in SCKAN is assigned a unique URI and is accompanied by a set of metadata annotations. These include human-readable labels from the curators, curatorial notes, and provenance information, with direct links to the source articles or artifacts from which curators extracted the connectivity information. Each class representing a neuron population in SCKAN is specified as equivalent to an intersection of a set of phenotypic relations defined using OWL class expressions. For instance, the class ‘Prostate:24,’ which denotes the neuron population ‘L2-L5 spinal cord to inferior mesenteric ganglion via lumbar splanchnic nerve,’ is defined as equivalent to the OWL expression shown in Listing 1, written in Manchester OWL syntax (Horridge et al., 2006).

**Listing 1.**
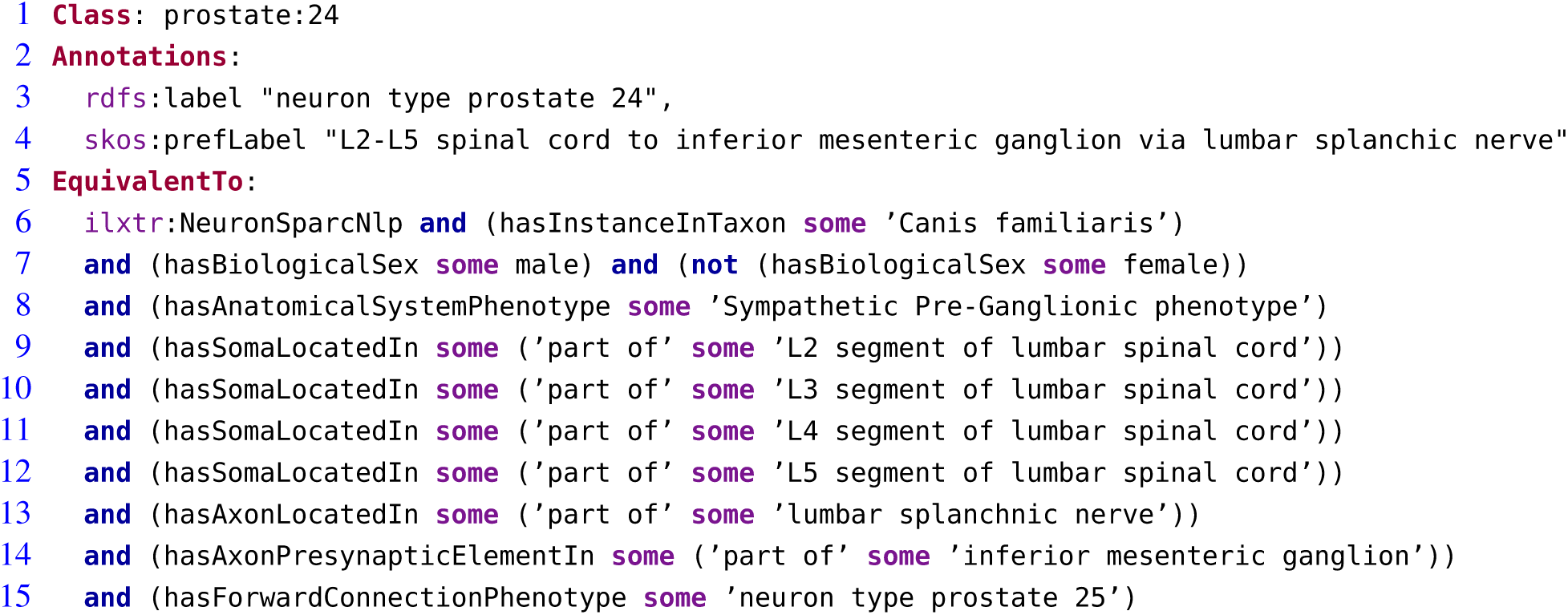
OWL Annotations and Equivalent Class Expression for a SCKAN neuron population

While the OWL axioms defining neuron populations in SCKAN can specify the locational context for each distinct neural segment, they are insufficient to capture the exact routing of the axonal course, particularly when multiple intermediate structures are involved. To address this, we augment the OWL axioms with an RDF list structure for each neuron population. Using an RDF list, we represent the axonal projection path by encoding the sequence of anatomical locations that an axon traverses as a hierarchical, ordered structure.

This approach captures the partial-order relationships among the pathway segments while preserving the overall order from origin to destination. We use a special property, `ilxtr:neuronPartialOrder`, to associate a neuron population with its partial-order tree. This approach also supports nesting ordered lists to represent complex axonal pathways that involve branching or convergence.

The code segment in Listing 2 provides an example of a nested RDF list structure to represent the axonal pathways of a SCKAN neuron population (prostate:24) in Turtle format. The hierarchical structures encoded as RDF lists can be queried using SPARQL to generate the corresponding adjacency list (Singh and Sharma, 2012), which represents pairwise relationships between consecutive locations along the pathway. The adjacency lists can then be utilized to render pathway diagrams using graph visualization tools and libraries.

**Listing 2.**
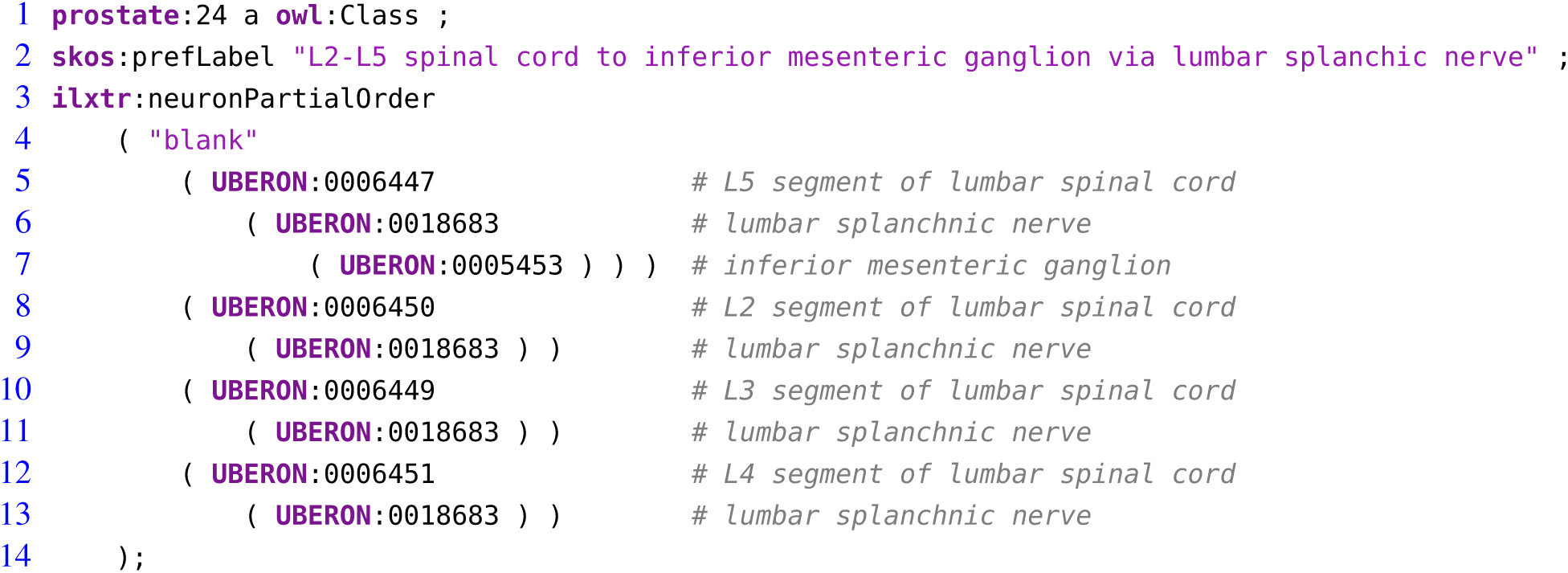
An example of RDF list representation specifying the axonal projection path for a SCKAN neuron population. Each node (element) in the RDF list corresponds to an anatomical structure.

The expert-contributed models in SCKAN, originally represented using ApiNATOMY (Kokash and de Bono, 2021), provide a representation significantly more expressive than NPO for describing physical anatomy and the detailed routing of neural pathways within larger tissue units (de Bono et al., 2022). In the context of neurons, this expressiveness enables the attachment of additional information to individual axons and dendrites, albeit with a concomitant increase in the complexity of the knowledge graph. Achieving similar granularity with NPO would require explicitly modeling axon and dendrite classes—a complexity deliberately avoided to simplify queries. Apart from the partial-ordering information for neural pathways, the additional anatomical detail and adjacency knowledge provided by ApiNATOMY were not incorporated into SCKAN. Each expert-contributed model represented in ApiNATOMY was carefully transformed into an NPO-based representation using the SCKAN-specific subset of properties shown in Figure 5. SCKAN currently provides a unified representation of connectivity for both ApiNATOMY-based expert-contributed models and literature-extracted connections, enabling a consistent approach to querying connections from both sources.

#### 2.4.1 Modeling Species, Sex, and Laterality

SCKAN facilitates the modeling of species and sex specificity for the animal in which a given neuronal connection was observed. Species-specificity in SCKAN is explicitly defined only when supported by at least one piece of experimental evidence documented in the published literature. Importantly, SCKAN operates under the open-world assumption for its reasoning, meaning that the absence of information does not imply nonexistence unless explicitly specified as such. Thus, species-specificity in SCKAN is represented through axioms of the form: ‘Population-X hasInstanceInTaxon some Species-Y’. This axiom indicates that the connection represented by ‘Population-X’ has been observed in ‘Species-Y’, while its presence or absence in other species is unknown. For example, Line 6 in Listing 1 specifies that the connections represented by ‘prostate:24’ have been observed in dogs. While it is possible that these connections may also exist in other species, such as humans, SCKAN leaves this possibility open, pending further evidence. While SCKAN adheres to the open-world assumption to remain faithful to available evidence, it also allows for the specification of negation axioms. For instance, a connection observed in one species could be explicitly negated for another if supported by evidence. However, as of now, no such negations have been introduced into SCKAN.

As discussed previously, SCKAN focuses exclusively on mammalian ANS connectivity. Consequently, connections in SCKAN that lack a species-specific axiom should, by default, be interpreted as mammalian. This default assumption does not imply that the connections are present across all mammals but rather that it has been observed in at least one mammalian species, with the exact species unspecified. All literature-extracted connections in SCKAN are associated with species-specific axioms. However, apart from the models of heart and bladder innervation, the remaining expert-contributed ApINATOMY models (as described in Table 1) currently lack species specificity and should therefore be considered mammalian by default. Species specificity for these models will be added in future releases as relevant data from published literature becomes available.

SCKAN represents sex specificity using axioms of the form: ‘Neuron-X hasBiologicalSex some Sex-Y’. If no sex is explicitly specified, the projection spanning the anatomical regions is assumed to exist in both males and females. Conversely, when a specific sex is explicitly specified, it signifies a sex difference. In such cases, SCKAN enforces a closed-world assumption by adding an explicit negation axiom for the other sex (see Line 12 in Listing 1 for an example).

Scientists are uncovering laterality differences in connections within certain species, where asymmetry exists between the left and right sides of the body. Such laterality differences are not encoded through logical axioms in SCKAN but are instead represented by assigning explicit laterality labels to the anatomical structures themselves. If a connection only exists on one side, e.g., in the sympathetic ovarian innervation via the superior ovarian nerve in the rat which innervates only the left ovary (Morán et al., 2009), a negation of the connection on the opposite side can be added to the SCKAN knowledge model. However, this form of modeling has not yet been implemented in SCKAN. Currently, we address such cases by providing a textual ‘alert note’ as an annotation.

#### 2.4.2 FAIR Vocabulary Management

All connections in SCKAN are annotated with the SPARC vocabulary (Surles-Zeigler et al., 2022) to ensure anatomical structures are used consistently across all SPARC systems. The overall strategy for the SPARC vocabulary is to utilize anatomical ontologies already in use by the community to conform to the FAIR principles and to ensure that SCKAN is interoperable with other projects in the Common Fund Data Ecosystem (CFDE, NIH Common Fund, 2021). As SCKAN contains connectivity information derived from multiple species, the pan-species anatomy ontology UBERON (Mungall et al., 2012) forms the “backbone” of its anatomical terminology efforts. Species-specific ontologies such as FMA (Rosse and Mejino Jr, 2003) and EMAPA (Hayamizu et al., 2013) are used to supplement as necessary. While SPARC is committed to reusing anatomical terms from standard community ontologies, we often require additional terms not present in community ontologies. To quickly mint identifiers for new terms, we utilize Interlex, an on-line vocabulary management system (Surles-Zeigler et al., 2022). To ensure the validity of new terms and that they are clearly defined, all proposed anatomical terms for SPARC go through specialized pipelines and a curatorial review before being added to the SPARC vocabulary. To make it easier to manage identifiers in SCKAN and perform QC, we restrict IDs for anatomical structures to UBERON and Interlex (ILX) IDs. If a term from another ontology such as FMA is required, it is imported into Interlex and an ILX ID mapped to the source ontology ID automatically generated.

It is critical that all new anatomical terms include the necessary semantics to support the queries and linkages required for effective use of SCKAN. All anatomical structures added to InterLex are explicitly related to UBERON terms, enabling combined reasoning over UBERON and the newly added terms. New anatomical structures are organized as children of anatomical entity (UBERON:0001096). Nerves and nerve plexi are added as children of nerve (UBERON:0001021), while specific ganglia are added as children of ganglion (UBERON:0000045). Anatomical structures may also be related to additional UBERON entities through the ‘part of’ relationship, as needed, to correctly associate individual connections with higher-order anatomy. In total, 328 anatomical terms were added to InterLex for SCKAN. The current version of SCKAN includes 242 distinct anatomical structures sourced from UBERON. The InterLex terms representing general anatomical structures are contributed back to community ontologies like UBERON to enhance their coverage of the central and peripheral nervous systems. We currently plan to submit 232 InterLex terms to UBERON.

#### 2.4.3 Simplified Representation of SCKAN

As described earlier, SCKAN is formally represented using OWL-DL, which supports the crucial requirements of automated reasoning for its formal knowledge statements. However, SCKAN’s OWL-based representation presents practical challenges in providing a flexible query mechanism for a broader audience unfamiliar with ontologies and formal knowledge modeling. While standard query languages like SPARQL and CYPHER are well-suited for writing data or instance-level queries, they become extremely difficult and tedious to use when crafting class-level queries involving complex ontological axioms. Retrieving high-level connectivity information from SCKAN’s OWL-based axioms requires writing complex, verbose queries that are only comprehensible to expert knowledge engineers trained in formal modeling. To mitigate this challenge, we developed an approach to generate a simplified representation of SCKAN, resulting in a representation we call Simple SCKAN.

Simple SCKAN representation provides an intuitive, higher-level abstraction over SCKAN’s formal axioms, enabling users to write streamlined queries without needing deep knowledge of OWL-DL jargon. It encapsulates SCKAN’s complex axioms by establishing a set of ‘shortcut relations’. Using SPARQL CONSTRUCT queries, we transform the intricate OWL-based statements in SCKAN into simpler RDF graph patterns for Simple SCKAN representation. After each SCKAN release, a Python script automates the transformation process. Simple SCKAN serves as a query-friendly layer on top of SCKAN. To query SCKAN, users need only familiarize themselves with the relational predicates supported by Simple SCKAN (Link to the documentation). Additional details and example queries can be found as part of the Simple SCKAN documentation.

The code segments in 3 and 4 provide a comparative view of querying SCKAN with and without Simple SCKAN representation. Both queries return the same results for “Give me the list of neuron populations in SCKAN along with their origins and destinations.” However, comparing the queries clearly demonstrates how SPARQL queries for SCKAN can become dramatically simpler to write using Simple SCKAN representation. The 18 lines of code, including complex OWL-specific jargon, in 3 (Lines 5-22) can effectively be replaced by 3 lines of simple code in 4 (Lines 4-7) using Simple SCKAN. The example also demonstrates how the Simple SCKAN version of the query is easier to comprehend and more manageable to write, test, and edit.

**Listing 3.**
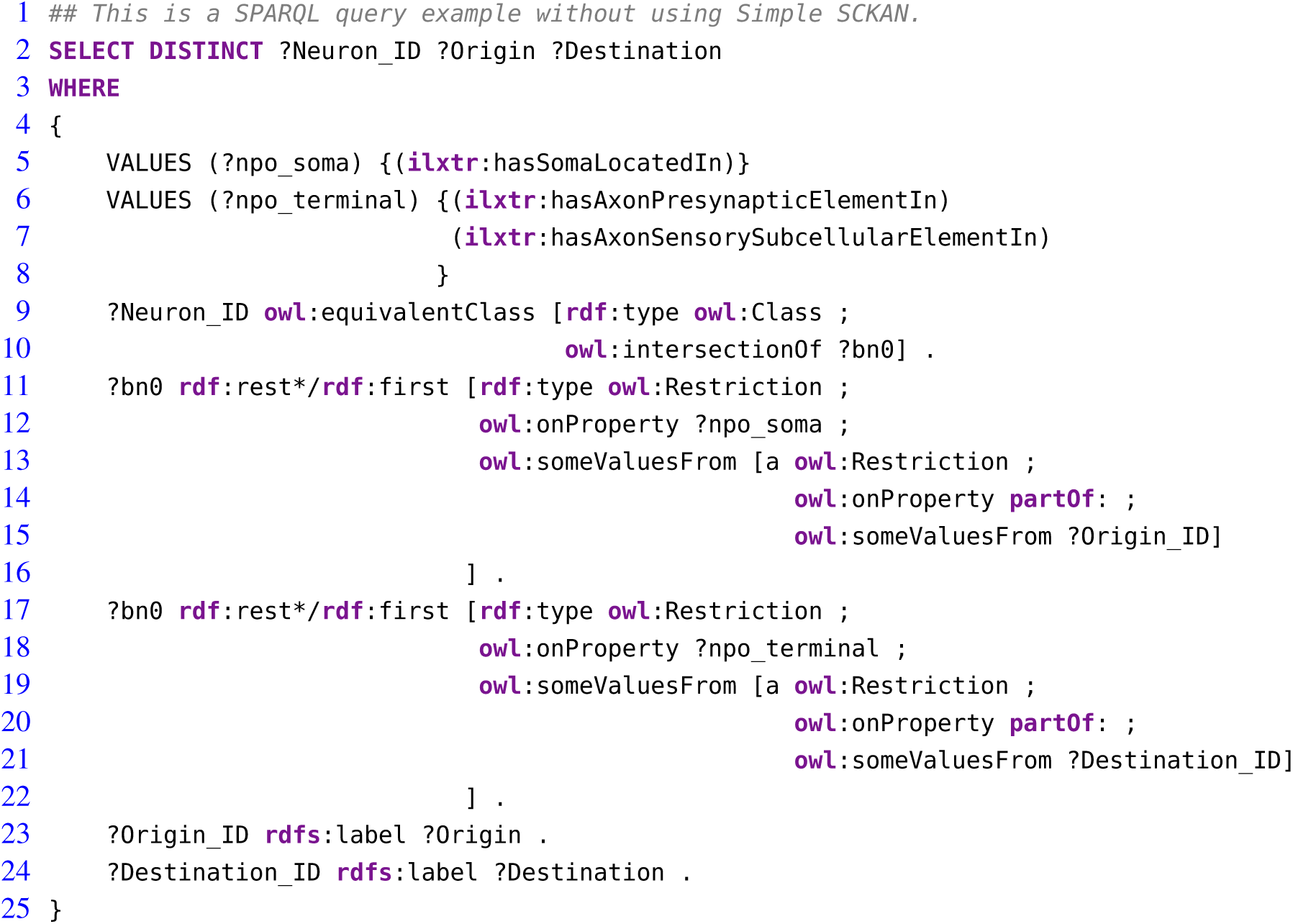
An example SPARQL query without using Simple SCKAN.

**Listing 4.**
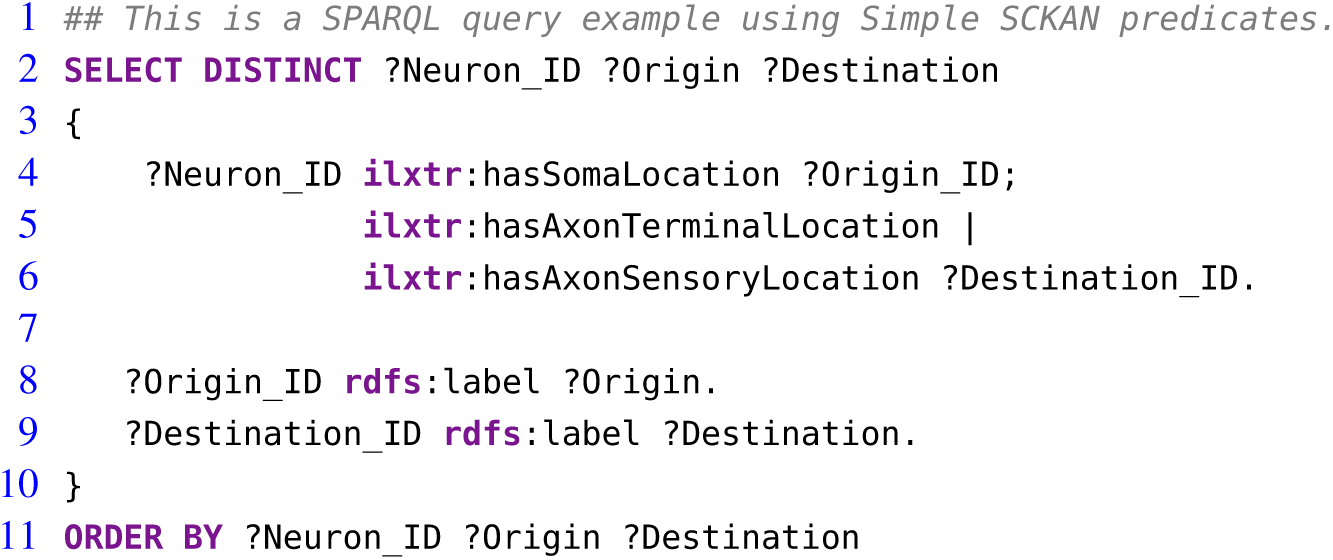
An example SPARQL query using Simple SCKAN predicates.

### 2.5 Accessing the Knowledgebase: SCKAN Endpoints

SCKAN is accessible through three different graph database endpoints namely: SciGraph (RRID:SCR 017576), Stardog (RRID:SCR 021223), and Blazegraph (RRID:SCR 026087), each serving distinct purposes (Figure 4). SCKAN is periodically loaded into SciGraph, a Neo4j-based graph database that hosts the SPARC knowledge graph. The SPARC knowledge graph leverages SCKAN to enhance the semantics of SPARC data, models, maps, and simulations with ANS connectivity for the SPARC Portal. Accessing SCKAN via SciCrunch also enables users to write queries using Cypher, a declarative query language for Neo4j databases. Some of the example use cases for the SciGraph endpoint are querying over the ApiNATOMY representation and providing terminology services for anatomical annotation.

Additionally, SCKAN is available through Stardog, a high-performance graph database equipped with a web interface called Stardog Studio. This interface streamlines the process of testing and editing SPARQL queries. We use Stardog primarily for its web interface, which allows storing and managing frequently used SPARQL queries for SCKAN. It provides an easy access for the knowledge engineers to check the contents of the knowledge base and share their queries with other user groups to expedite their interaction with SCKAN.

Lastly, SCKAN can be accessed through a public Blazegraph endpoint that does not require registration or authentication. Blazegraph is an ultra-high-performance graph database that supports RDF and SPARQL APIs. We utilize Blazegraph for its rapid response times in executing complex SPARQL queries, which may otherwise take several minutes with other graph database systems. Simple SCKAN representation of SCKAN is available via the Stardog and Blazegraph endpoints; however, it is not accessible via the SciGraph endpoint. Links for Accessing SCKAN:

*•* SciGraph Endpoint: Link to the GitBook document
*•* Stardog Endpoint: Link to the GitBook document
*•* Jupyter Notebook on SCKAN Example Queries: Link to the GitHub document

#### 2.5.1 SCKAN Explorer: A Query Interface for SCKAN

To streamline the process of extracting basic connectivity information from SCKAN, we developed SCKAN Explorer (RRID:SCR 026080, https://services.scicrunch.io/sckan/explorer/)—an intuitive query interface specifically designed for SCKAN’s knowledge curators. It serves as a prototype interface to support basic curatorial tasks, enabling users to quickly check and verify existing connections in SCKAN without the technical hurdle of writing SPARQL queries. SCKAN Explorer facilitates streamlined data input by providing auto-complete suggestions for SCKAN-specific anatomical locations and their synonyms. It provides filtered search results based on origin (soma location), destination (axon terminal or sensory terminal location), and/or via location (axon pathways such as nerves, nerve plexuses, or nerve fibers). Additional search criteria include connection phenotype (e.g., sympathetic, parasympathetic), named connectivity models, neuron ID, species, and target organ. SCKAN Explorer also enables the visualization of connectivity pathways for individual neuron populations, including their synapses.

Figure 6 provides example screenshots from SCKAN Explorer. In this example, selecting ‘ANS:Sympathetic Pre-Ganglionic’ as the connection phenotype for the Female Reproductive System returns 10 neuron populations from SCKAN. The screenshot at the bottom-left displays an example population (femrep:30) along with its phenotypic properties, including species and sex specificity, links to the references from which the connectivity information was derived, and a tabular representation of its pathways between the origin and destinations. On the right, the pathway diagram for the population is shown, including its synaptic forward connections. Exploring the connections in this way helps curators verify the accuracy and completeness of their added information. For instance, the ‘End Organ’ query traverses the partonomy chain of major organ systems to retrieve all populations that terminate in one or more subregions of a selected organ. Comparing the number of populations returned can reveal gaps in the partonomy chains in UBERON or new terms added via InterLex. When such issues are identified, they are resolved by adding the missing partonomy chain to InterLex and importing it into SCKAN for the next release candidate.

**Figure 6.**
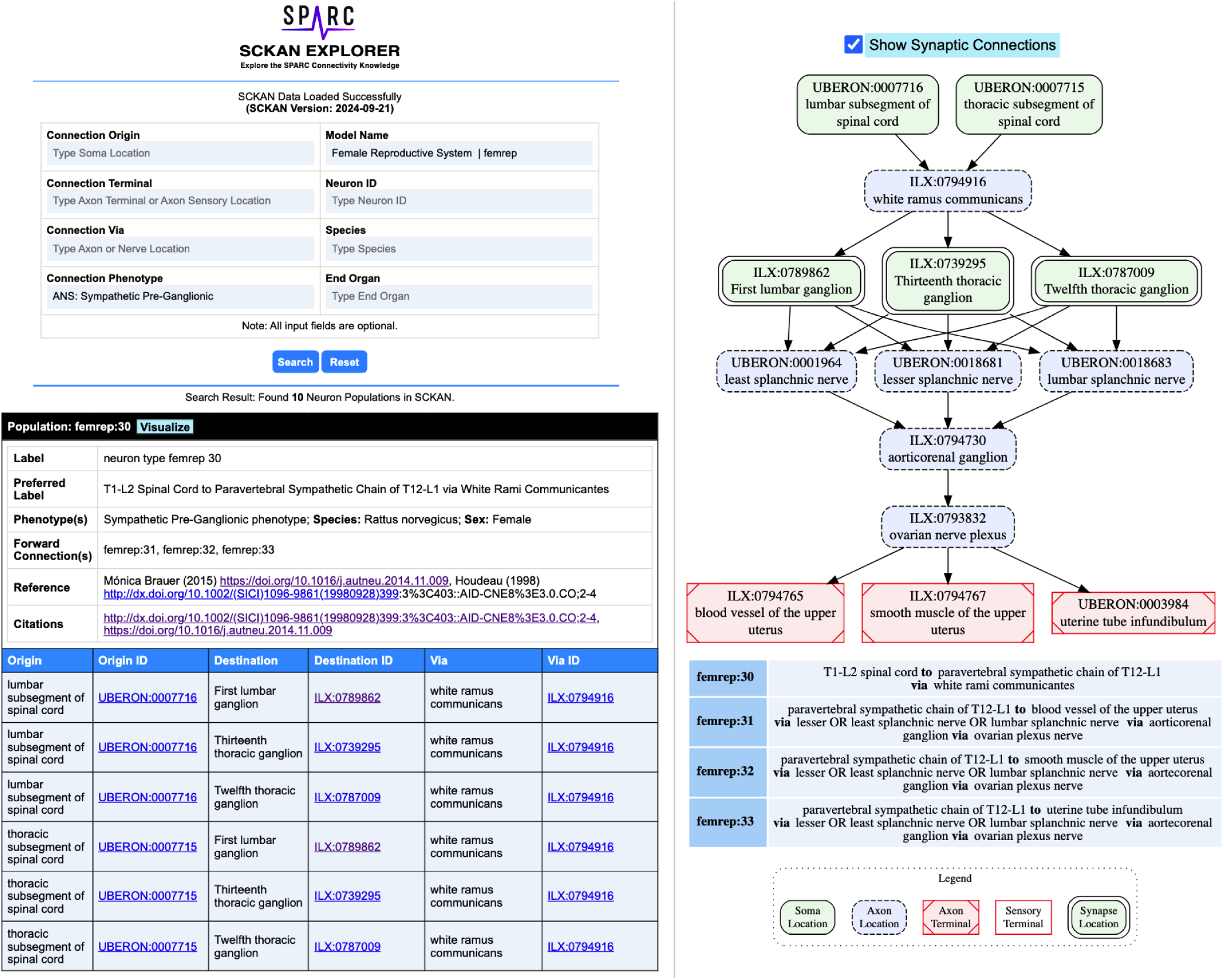
Example screenshots from the SCKAN Explorer interface (https://services.scicrunch.io/sckan/ explorer/).

## 3 RESULTS

### 3.1 SCKAN Contents: Statistical Summary

SCKAN releases occur on a roughly quarterly basis. The current SCKAN release (Version: 2024-09-21) represents the end of the initial phase of content driven by the SPARC project, covering ANS connectivity for most major organ systems. It includes some sensory and motor connections as well (see Table 3), especially those related to the vagal system, but these were not the focus of this version of SCKAN. In this section, we provide an overview of SCKAN’s content coverage.

Table 2 summarizes the current coverage of SCKAN, categorized by major organs and systems. As shown, SCKAN includes 112 expert-contributed populations and 180 literature-extracted populations. The table also reports the count of distinct locational phenotypes associated with each organ or system category. Notably, SCKAN has limited coverage of sensory terminals, as sensory projections were not within the scope of SPARC.

**Table 2.** Coverage of SCKAN in terms of its neuron population counts categorized by different organs and systems, along with the counts of specific locational phenotype for each category.

| Population Category | Population Count | Soma Location | Axon Location | Axon Terminal | Sensory Terminal |
| --- | --- | --- | --- | --- | --- |
| **Expert-Contributed Models** |  |  |  |  |  |
| Bolser-Lewis Model of Defensive Breathing | 29 | 9 | 37 | 15 | 5 |
| Keast Model of Bladder Innervation | 20 | 27 | 57 | 20 | 5 |
| SAWG Model of Bronchomotor Control | 6 | 19 | 11 | 9 | 0 |
| SAWG Model of the Descending Colon | 18 | 11 | 17 | 20 | 4 |
| SAWG Model of the Pancreas | 5 | 11 | 19 | 5 | 0 |
| SAWG Model of the Spleen | 5 | 8 | 10 | 5 | 1 |
| SAWG Model of the Stomach | 10 | 6 | 8 | 8 | 6 |
| UCLA Model of the Heart | 19 | 33 | 35 | 19 | 7 |
| **Total** | **112** |  |  |  |  |
| **Literature-Extracted Connections** |  |  |  |  |  |
| Cranial Nerve Connections | 10 | 6 | 12 | 10 | 0 |
| Female Reproductive System | 60 | 39 | 26 | 54 | 7 |
| Kidney Connections | 27 | 53 | 44 | 50 | 9 |
| Liver Connections | 26 | 22 | 20 | 27 | 4 |
| Male Reproductive System | 21 | 29 | 9 | 8 | 0 |
| Sweat Gland Connections | 8 | 28 | 15 | 18 | 0 |
| Uncategorized Connections | 28 | 39 | 44 | 25 | 2 |
| **Total** | **180** |  |  |  |  |

Table 3 summarizes the counts of SCKAN populations based on specific phenotypes, categorized by the phenotypic dimensions of species, sex, locational phenotypes, functional circuit role, circuit role, and ANS phenotype. As outlined earlier, SCKAN represents species-specificity through explicit axioms, while connections without such axioms are assumed to be mammalian by default. As shown in Table 3, SCKAN currently contains 88 neuron populations with ”unassigned” species. Following the default assumption stated in Section 2.4.1, these populations are assumed to exist in at least one unspecified mammalian species.

**Table 3.** Counts of SCKAN neuron populations based on different phenotypic dimensions.

| Dimension | Phenotypic Value | Count |
| --- | --- | --- |
| Species | Cat | 5 |
|  | Dog | 22 |
|  | Guinea Pig | 1 |
|  | Human | 99 |
|  | Mouse | 11 |
|  | Pig | 19 |
|  | Rat | 86 |
|  | Unassigned | 88 |
| Sex | Female | 69 |
|  | Male | 25 |
| Locational Phenotype | Soma Location | 292 |
|  | Dendrite Location | 42 |
|  | Sensory Axon | 37 |
|  | Axon Terminal | 290 |
|  | Sensory Terminal | 42 |
|  | Axon Location | 256 |
| Functional Circuit Role | Excitatory | 4 |
|  | Inhibitory | 4 |
| Circuit Role | Intrinsic | 4 |
|  | Motor | 9 |
|  | Sensory | 40 |
| ANS Phenotype | Enteric | 1 |
|  | Parasympathetic | 77 |
|  | Sympathetic | 131 |

In contrast, Table 2 also lists the counts of populations associated with specific species, such as the 99 neuron populations tied to humans and the 86 tied to rats. It is important to note that, under SCKAN’s open-world assumption, observing a connection in one species does not imply its presence or absence in others. Thus, the species-specific counts reported in the table should not be considered mutually exclusive. For example, the 18 neuron populations in the UCLA Model of the Heart were observed to be common across dogs, pigs, and humans. Currently, SCKAN includes 69 populations associated with female anatomy and 25 with male anatomy, as shown in Table 3.

The sex-specific populations in SCKAN are accompanied by explicit negation axioms, as outlined in Section 2.4.1, making them mutually exclusive to the opposite sex (e.g., the connections represented by the female-specific populations cannot exist in males). The remaining 198 populations, lacking sex-specific axioms, are assumed to exist in both sexes, as noted in Section 2.4.1.

As previously mentioned, the current focus of SCKAN is on neural connectivity associated with the distinct subdivisions of the ANS. Consequently, the number of populations associated with ANS subdivision phenotypes predominates the current knowledgebase as reported in Table 3. Nevertheless, we do have small numbers of sensory and motor neuron populations in SCKAN, largely due to their inclusion in some of the connectivity models provided by experts. In addition, as the vagus is a major target of SPARC efforts, we include sensory populations arising from the nodose ganglion, a sensory ganglion that is physically part of the vagus nerve.

### 3.2 Competency Queries

SCKAN was designed to answer questions relevant to ANS connectivity. To demonstrate the functionality of SCKAN we developed a set of competency queries (CQ). In this section, we work through the CQs and examine the results. For each question, we provide the SPARQL query and a visualization of the query results. All of the SPARQL queries listed in this section are written using Simple SCKAN. Navigate to the SCKAN CQS Repository for the raw query results in CSV format, along with the visual diagrams produced by the SPARQL queries included in this section. For the first three CQs we used RAWGraphs

(RRID:SCR 026082) to produce the visual query results. RawGraphs (https://github.com/rawgraphs/ rawgraphs-app) is an open-source tool to create customized vector-based visualizations on top of *d*3*.js* library. For the fourth CQ, we developed a custom visualizer utilizing GraphViz (RRID:SCR 026086, https://graphviz.org/), an open-source graph visualization library to represent structural information as diagrams.

#### CQ-1: What connections terminate in the bladder? What are the origins of those connections and what are the exact parts of the organ the connections terminate. What nerves are involved in those connections?

**Listing 5.**
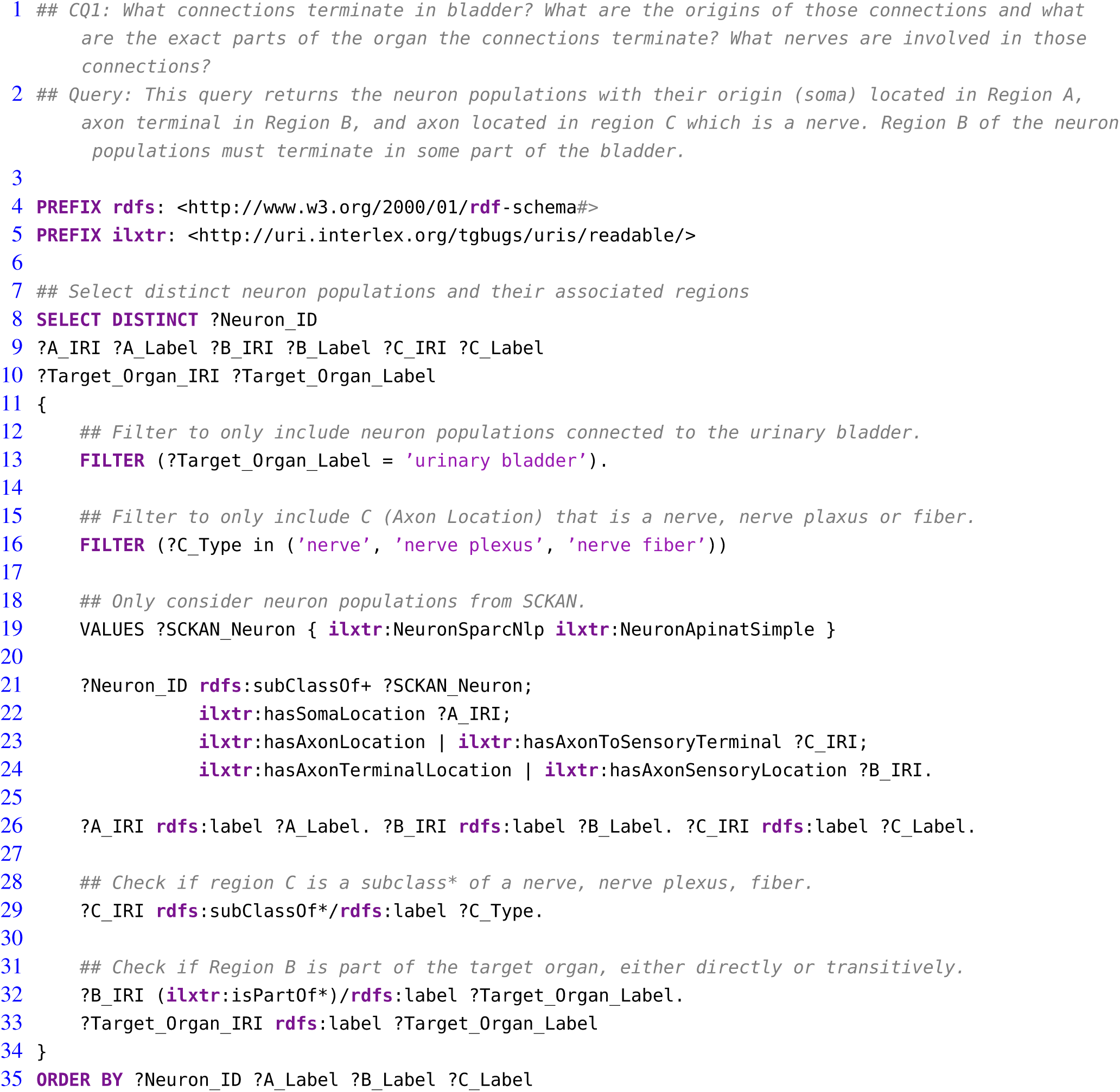
The SPARQL query for CQ1 to retrieve the connections projecting to the urinary bladder, detailing their origins (Region A), termination points within the bladder (Region B), and axon locations within associated nerves (Region C).

The SPARQL query (Listing 5) retrieves the required information based on filtering a set of neuron populations in SCKAN with specific connections to the urinary bladder. The query identifies populations with their soma located in one region (Region A), their axon terminal or axon sensory terminal located in another region (Region B), and the axon potentially passing through or being located in a third region (Region C) which must be a nerve. The focus of the query is on neuron populations where Region B must satisfy the constraint of being a part of the bladder. The query for all populations that terminate in the bladder returns connections contained within 6 distinct neuron populations that have axon terminals in some part of the bladder. The visual query results in Figure 7 shows the origins of these connections (right panel) and the nerves through which they traverse, including bladder nerve, pelvic splanchnic nerve, and hypogastric nerve.

**Figure 7.**
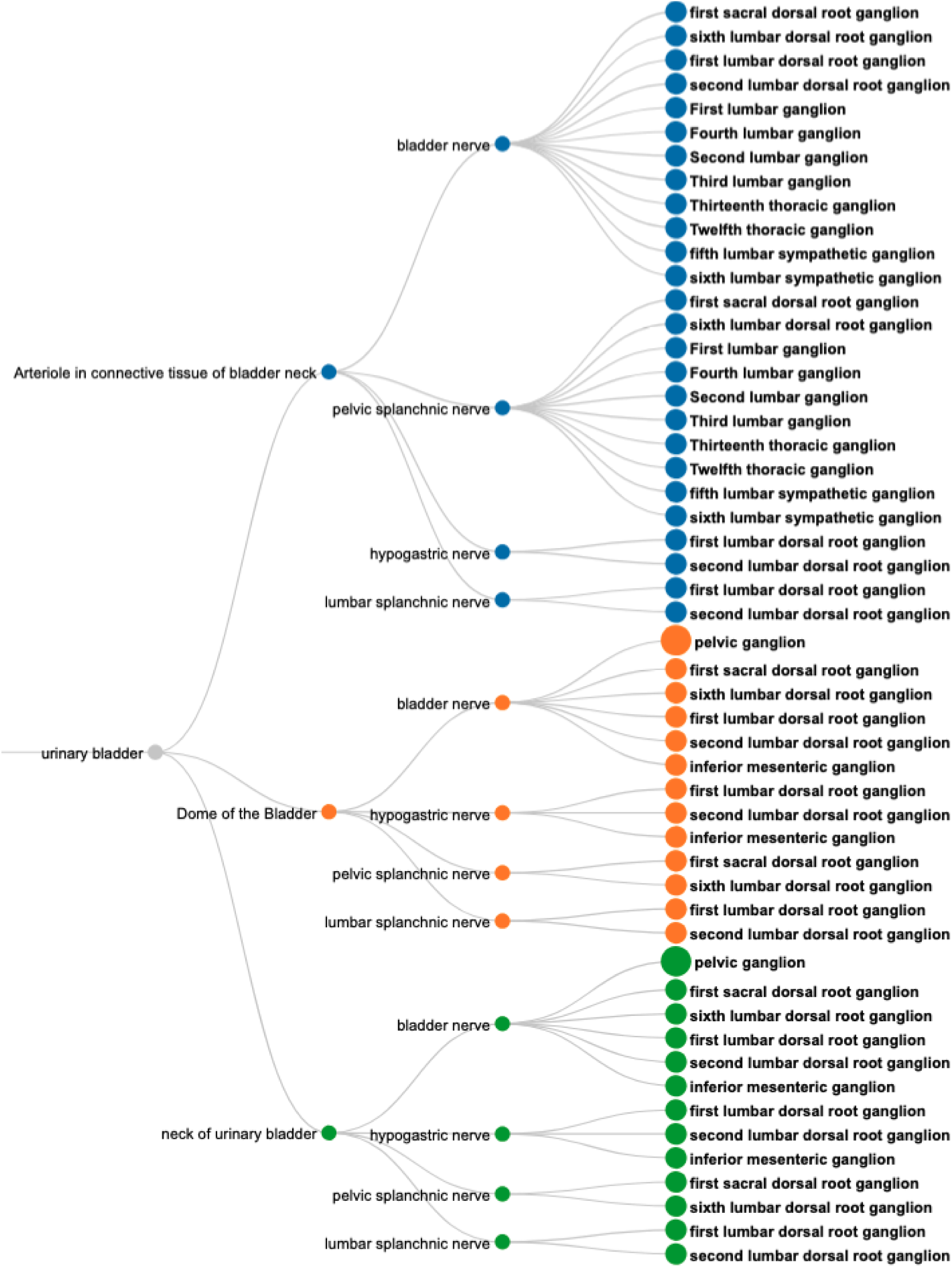
A visual query result for *CQ*1: Neural pathways terminating in the bladder, illustrating their origins (nodes on the right), nerve involvement (nodes in the middle), and specific bladder regions targeted (nodes on the left). Nodes are color-coded to distinguish the connections based on the terminal structures in the bladder.

#### CQ-2: What are all origins and destinations of neuron populations traveling through the vagus?

**Listing 6.**
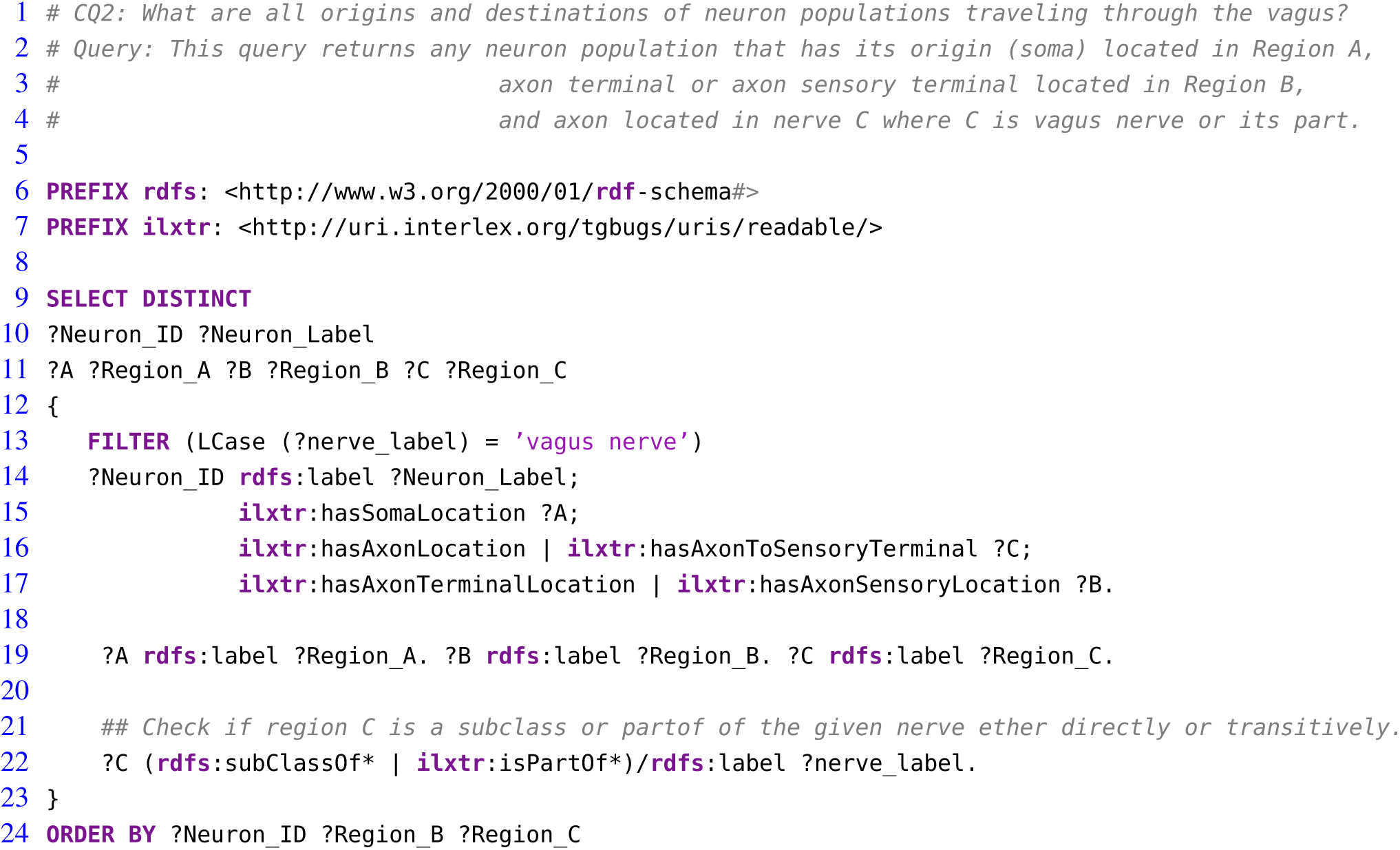
The SPARQL query for *CQ*2 to retrieve neural connections with origins (Region A) and destinations (Region B) that travel through the vagus nerve or its branches (Region C).

The SPARQL query (Listing 5) retrieves the required information based on filtering a set of neuron populations in SCKAN with specific connections via the vagus nerve or its branches. Figure 8 shows the visual results for a portion of connections arising from the inferior vagus X ganglion (nodose ganglion). Please navigate to the full visual result for the remaining connections.

**Figure 8.**
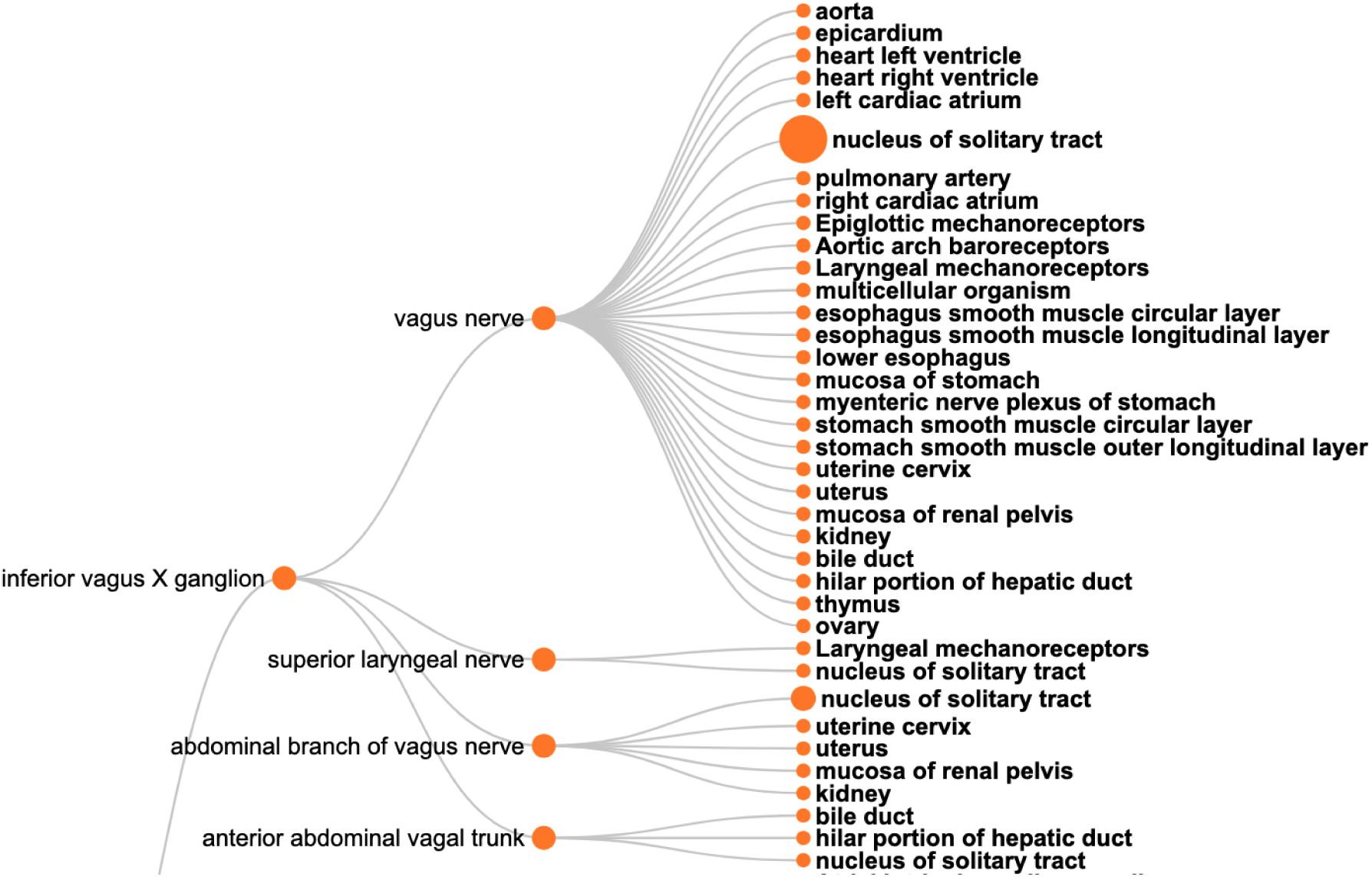
A visual query result for *CQ*2: The origins (the left nodes) and destinations (nodes on the right) of neural connections via the vagus nerve, highlighting its diverse innervation across multiple organs.

The query results show that SCKAN currently has 35 such neuron populations arising from the following 7 distinct origins: inferior vagus X ganglion (nodose ganglion), dorsal motor nucleus of vagus nerve, nucleus ambiguus, superior cervical ganglion, left nodose ganglion, right nodose ganglion, and the nucleus of solitary tract. The query returns 43 distinct destination structures across different organs innervated by the vagus nerve or its branches. The query recognizes the following partonomic vagus nerve branches from SCKAN: External and internal branches of superior laryngeal nerve, External and internal branches of inferior laryngeal nerve, superior laryngeal nerve, abdominal branch of vagus nerve, esophageal vagus trunk, vagus X nerve trunk, anterior and posterior abdominal vagal trunks, and the posterior hepatic branch of vagus nerve.

#### CQ3: What anatomical structures might be affected by perturbation of the inferior mesenteric ganglion?

This type of query represents one of the driving motivations for SPARC and SCKAN, the ability to better predict on and off target effects for a given electrode placement. Based on the anatomical information in SCKAN, we can pose this query by retrieving all populations that have any part (soma, axon, axon terminal) in the IMG and then retrieving their origins and destinations. The SPARQL query for CQ3 is provided in Listing 7. Figure 9 shows the 55 unique structures that could potentially be affected through a perturbation of the IMG.

**Listing 7.**
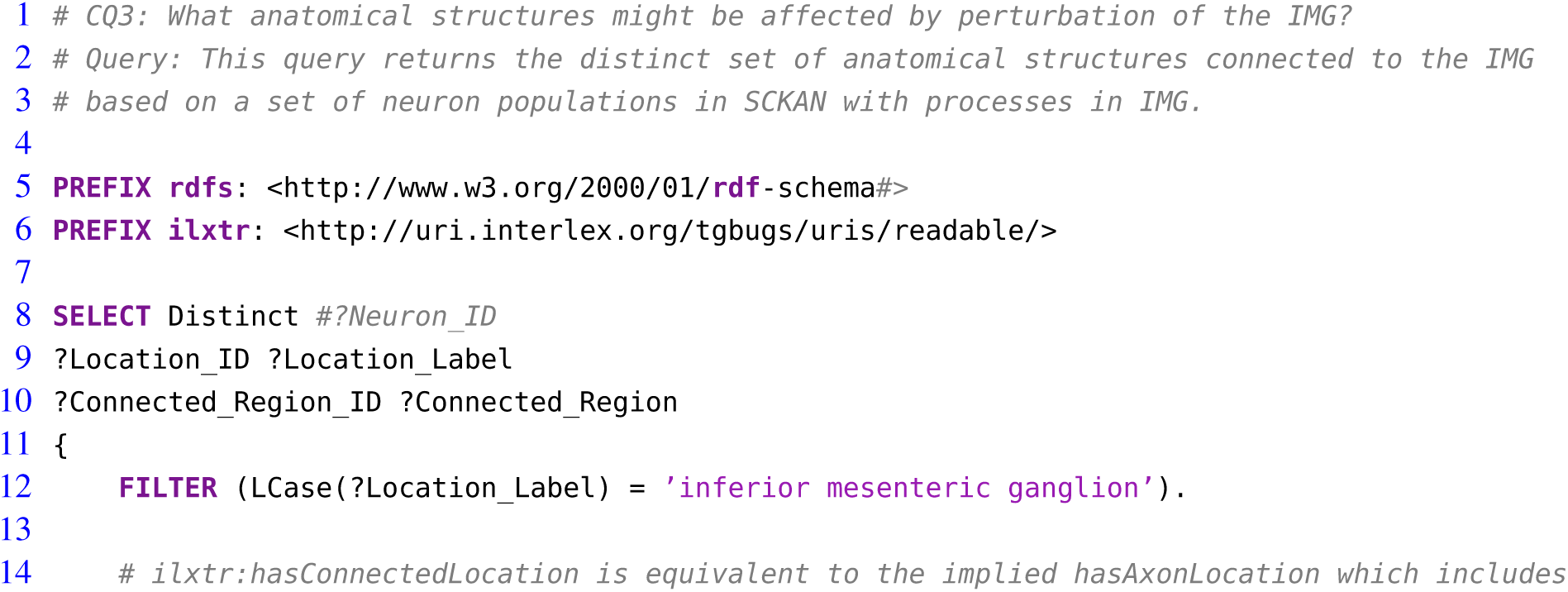
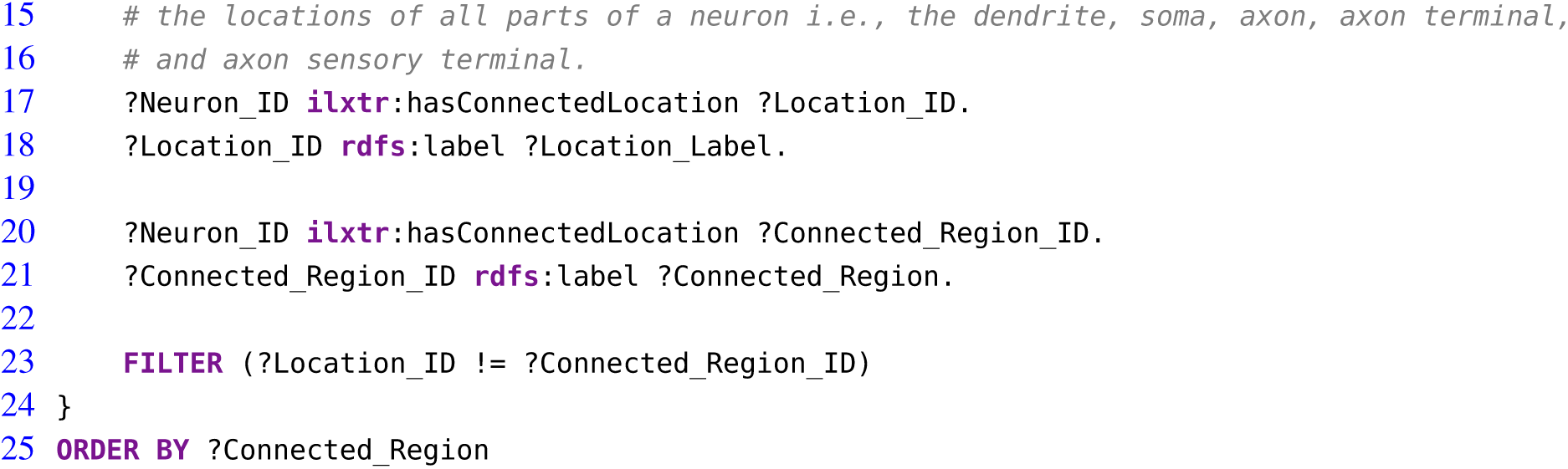
The SPARQL query for CQ3 to identify anatomical structures potentially affected by perturbations of the inferior mesenteric ganglion (IMG) based on neuron populations in SCKAN.

**Figure 9.**
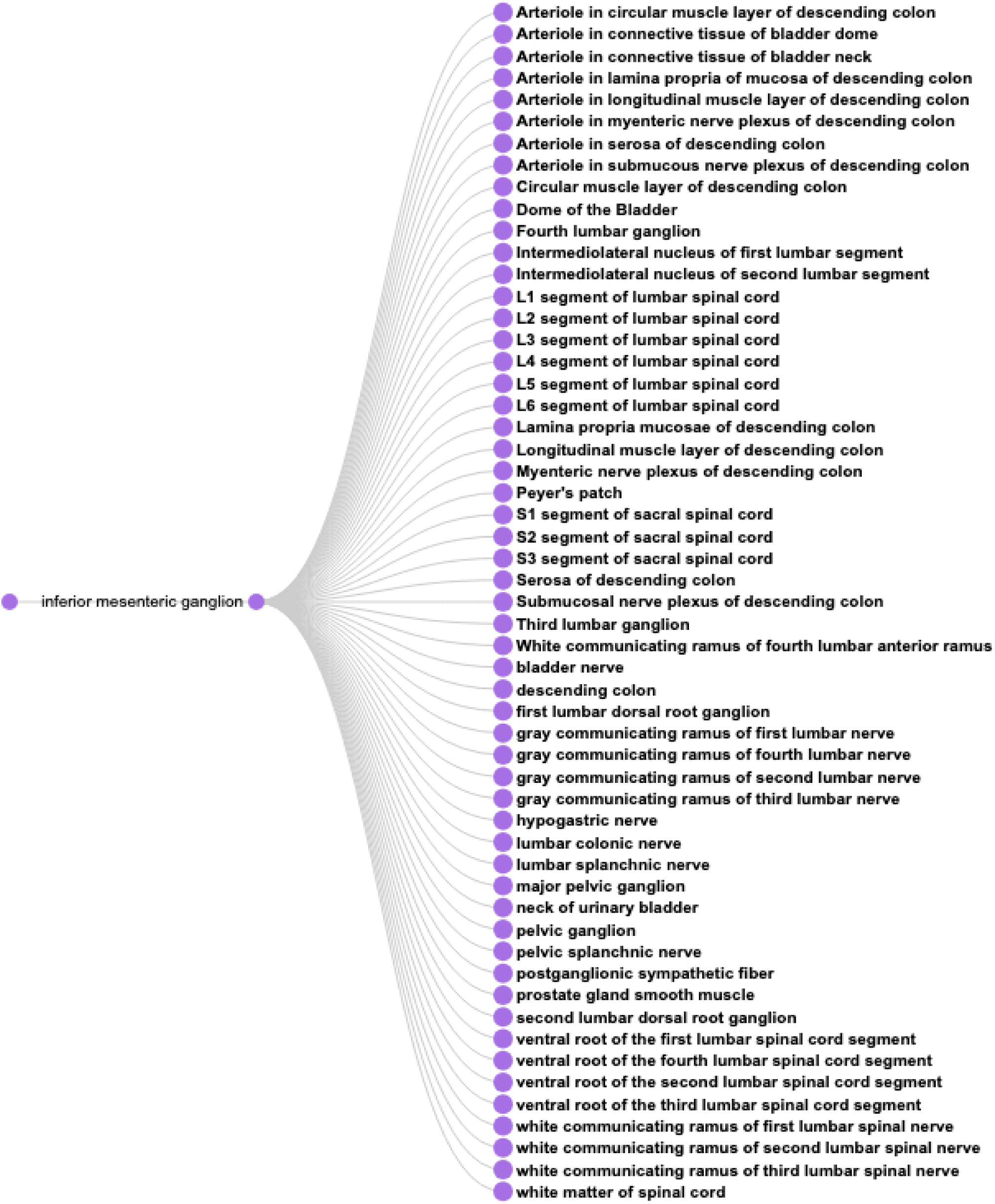
A visual query result for *CQ*3: The result shows the 55 potential anatomical structures in SCKAN that might be affected by perturbing the inferior mesenteric ganglion.

#### CQ4: How is sympathetic innervation supplied to the ovaries? Show the origins, terminations, and routing for both pre- and post-ganglionic connections

This query looks for all pre- and post-ganglionic sympathetic populations connected via a forward connection where the post- ganglionic neurons terminate in the ovary. Neither of the current GUI’s make it easy to answer this question. A query through SCKAN Explorer can return all post-ganglionics by entering “ovary” in end organ (both sympathetic and parasympathetic), but retrieving pre-ganglionics requires knowledge of the specific pre-ganglionic populations. SCKAN currently connects pre- and post- ganglionic populations through the ‘hasForwardConnection’ property, but the property is not reciprocal.

Listing 8 provides the SPARQL query for *CQ*4. The query result suggests the innervation involves 5 preganglionic and 6 postganglionic sympathetic neuron populations linked by the synaptic forward connections. Figure 10 shows the visualization of the result. The figure shows the origins, terminations, and routing for both pre- and post-ganglionic connections along with 5 synaptic locations. It should be noted that the visual query result is simplified for easier comprehension by collapsing the nodes representing multiple adjacent segmental structures into single ones (e.g., using a single node like ‘T10-L1 segments of spinal cord’ instead of having 4 different nodes representing each individual segment between T10 and L1). This is an example of a query where the raw query results cannot be comprehended effectively without the use of visual diagrams. Visualizing this complex query result demonstrates how SCKAN is laying the foundations for modeling and simulations of neuromodulation therapies. Note that the sympathetic innervation of the ovary is an example of an asymmetrical connection, where in the rat, innervation of the left ovary is different than the innervation of the right ovary (Morán et al., 2009).

**Figure 10.**
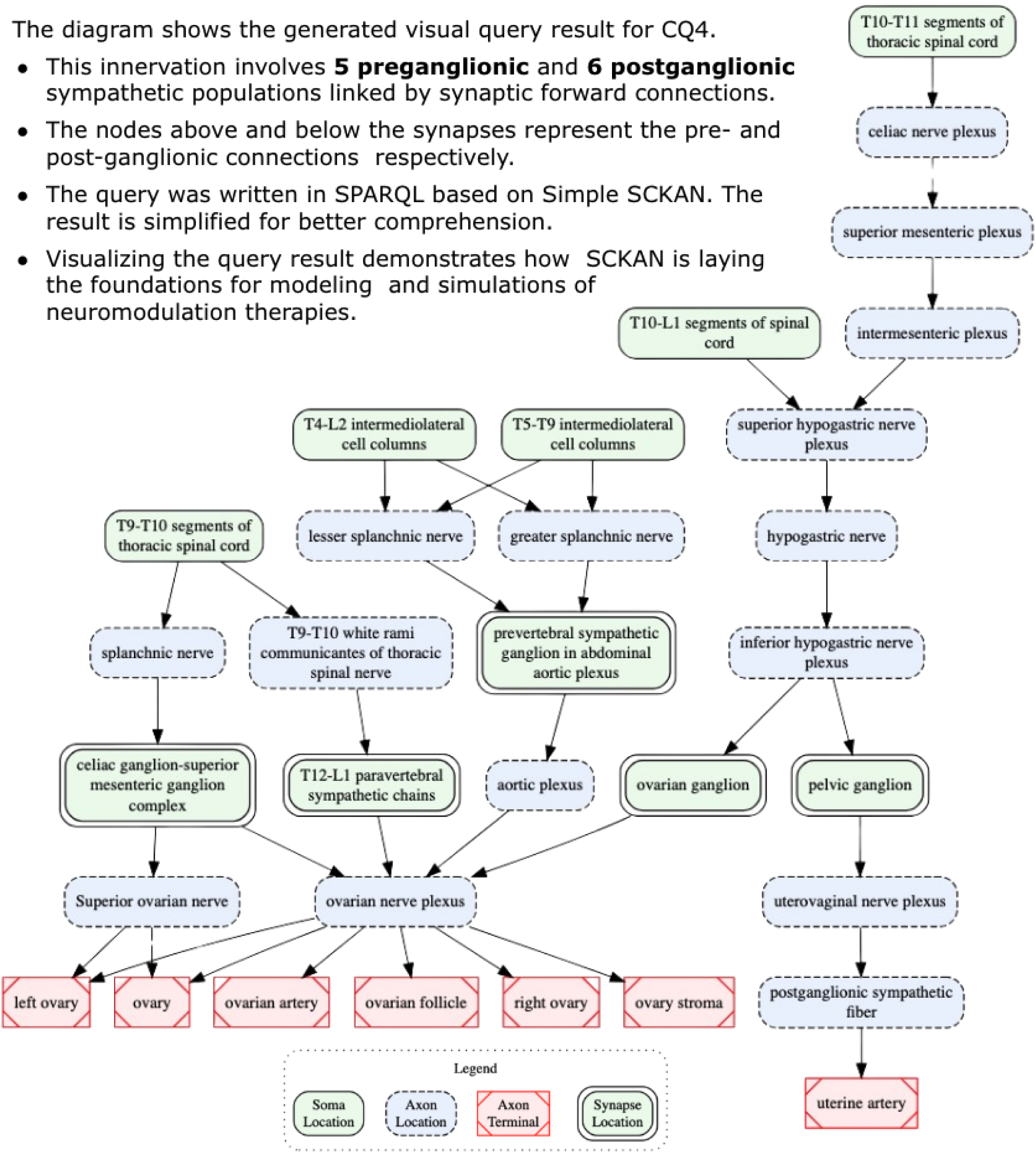
A visual query result for *CQ*4: detailed circuitry involved in sympathetic innervation of the ovaries, depicting pre- and post-ganglionic connections, routing, and synaptic locations.

**Listing 8.**
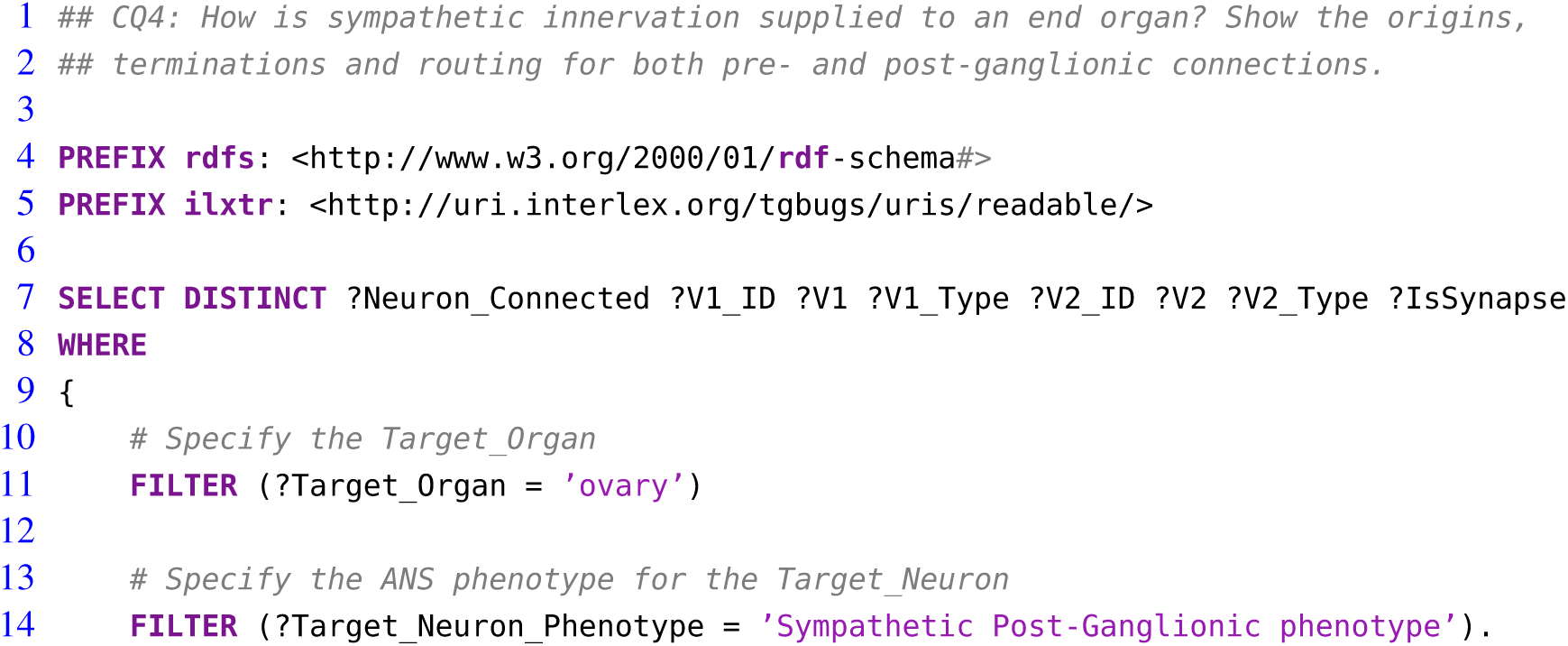
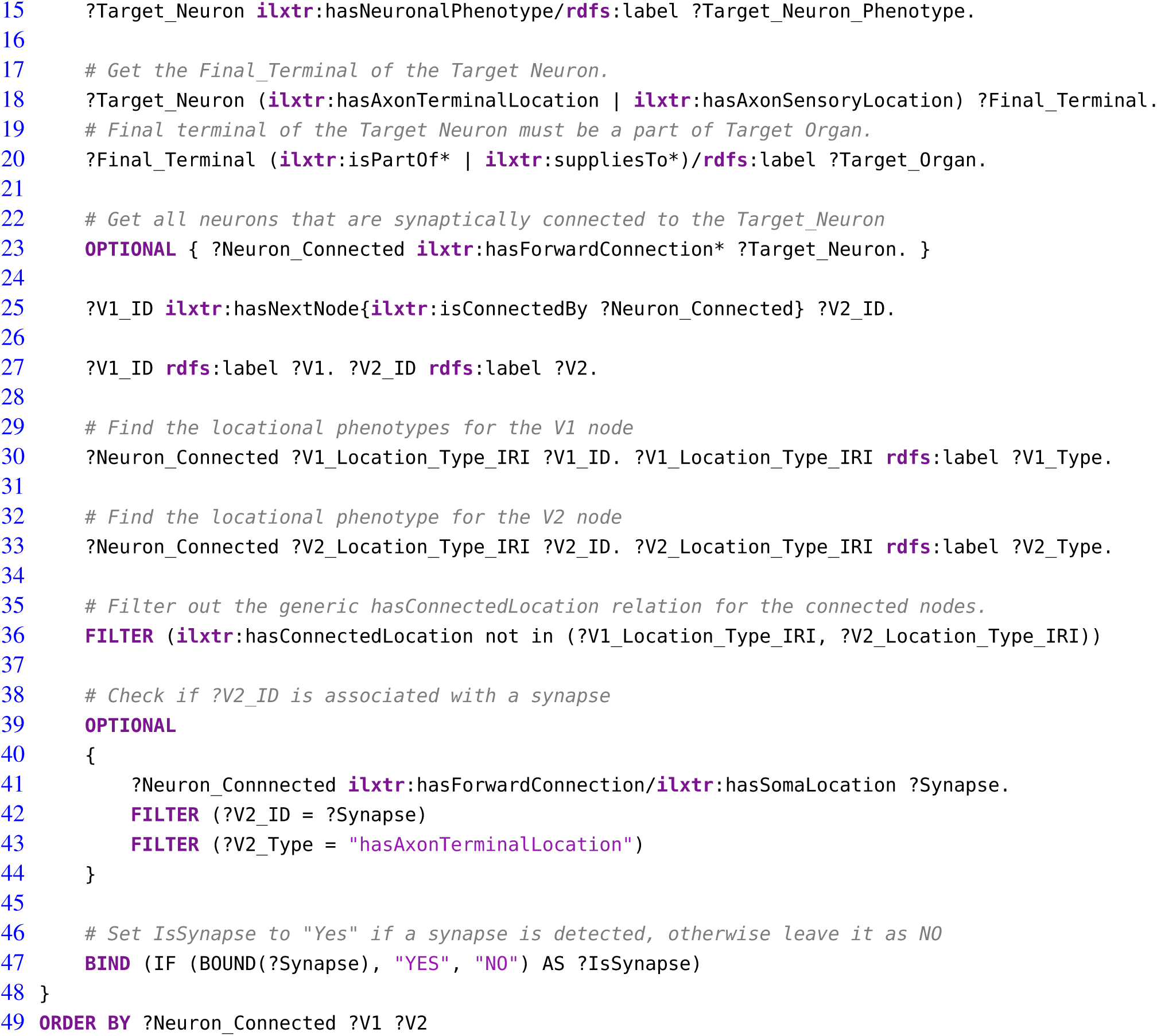
The SPARQL code for *CQ*4 to trace sympathetic innervation pathways to the ovary, detailing origins, terminations, and routing of pre- and post-ganglionic connections, including synapse locations.

## 4 DISCUSSION

Neuromodulation therapy and increased attention to the gut-brain axis are shining spotlights on the importance of understanding the anatomical, functional and molecular organization of the peripheral nervous system. To our knowledge, SCKAN represents one of the only connectivity knowledge bases that focuses on the peripheral nervous system rather than the central nervous system e.g., the Allen Mouse Brain Connectivity Atlas (RRID:SCR 008848, http://connectivity.brain-map.org/), BAMS Neuroanatomical Ontology (RRID:SCR 004616, https://bams1.org/ontology/viewer.php) (Bota and Swanson, 2008), and CoCoMac (RRID:SCR 007277, https://cocomac.g-node.org/). Although the current version of SCKAN contains mostly ANS connectivity, reflecting the priorities of the NIH SPARC Program, we are in the process of extending the content with peripheral sensory and motor pathways. The expansion reflects the concomitant expansion of the mission of the SPARC Portal and associated infrastructure as the NIH SPARC Program winds down.

SPARC recently became an open data platform for data, models and simulations involving the peripheral nervous system and its end organ, spinal cord, and brain stem interactions. In this way, SPARC complements the infrastructure established through the NIH BRAIN Initiative and similar projects around the world that focus on the CNS. We expect that SCKAN will provide a valuable source of connectivity information to complement the US BRAIN CONNECTS program that is getting underway.

SCKAN was designed to represent anatomical connections in a way that details routing information of axons and locations of somata, axon terminals and axon sensory terminals. In this version of SCKAN, we do not focus on detailed synaptic connections and so dendrites are largely not included. SCKAN connections are automatically visualized via the SPARC’s interactive 2D and 3D anatomical and functional maps (available at https://sparc.science/apps), which are updated with each release of SCKAN on a quarterly basis. The flat maps contain enough detail to understand basic pathways including nerves but retain a level of abstraction to make these routes easy to understand, much like the static images of ANS that are present in resources such as medical textbooks. SCKAN contains more detailed routings which can be returned via queries and visualized in detail. Because SPARC flatmaps are not static drawings but are generated via SCKAN, they can be kept up to date as new data are added or edited.

To ensure accuracy of the detailed connections in SCKAN, we developed a curatorial workflow that provided multiple checks on connectivity from the initial identification of connections to their coding and query in SCKAN. The information encoded in SCKAN is complex, with assignment of individual parts of a neuron to different locations. Often these different parts of a connection had to be pieced together from multiple scientific articles, especially figures. These parts must then be reassembled in the correct order to represent a connection. Keeping track of this information and allowing its progressive refinement through the pipeline was error prone and time consuming when using tools such as Google Sheets. For that reason, we developed a set of tools to support the curators to manage review of sentences extracted from the literature, distillation of knowledge from multiple sources into discrete neuron populations, expression of connections in terms of locations of soma, axon segments and axon terminals followed by mapping to UBERON or Interlex identifiers, and finally coding into OWL. These tools made it easier for domain experts, curators and knowledge engineers to work together with a shared understanding of what was to be encoded into SCKAN. They also allowed curators to query and visualize the contents of SCKAN for additional QC and determine whether connections mentioned in the literature were missing.

The curation pipeline involved 3 sets of skills: (1) a curator with neuroscience knowledge sufficient to read and interpret connectivity sources and basic understanding of the SCKAN data model and how to work with ontologies; (2) a domain expert with deep knowledge who could review the information for accuracy and recommend authoritative sources; (3) a knowledge engineer who could do the deep semantic encoding required. In our case, these skills were embodied in multiple individuals. The curator played the crucial go-between role of identifying populations through translation of the literature and expert knowledge and communicating it in a way that lent itself to knowledge engineering. Initially, we asked domain experts do this, but it was difficult for them to express the connections in a form that was easy for the knowledge engineers to encode.

Ontological representation is deterministic and not probabilistic, and therefore connectivity statements needed to be expressed in a way that lent themselves to encoding as a set of triples. This type of simplification is often difficult for researchers with deep knowledge of the nuances of connections. We found that constructing single, formulaic sentences in the form of *A* projects to *B* via *C*, each describing a single population, along with relevant parts of papers supporting the population, was the most efficient way for our experts to review them. These sentences also served the critical role as a check throughout the encoding process that what is being expressed in OWL reflects the connection accurately. The new Composer tool allows a curator either to construct a statement manually or automatically by adding the locations of cell soma, axons and axon terminals into a structured form.

### 4.1 Cell Populations Vs Cell Types

Unlike many connectivity databases, we elected to base connectivity on a neuronal model rather than a regional model. Connections are modeled via the locations of cell bodies, axons and axon terminals which reside in anatomical locations. Axon locations are modeled segmentally so that the routing of an individual axon, which may traverse many anatomical structures and travel via multiple nerves can be accurately traced. Although there is a computational cost to this approach, we elected to employ the neuron populations model for multiple reasons. First, we can more easily accommodate multi-scale connectivity into the same model in that we can provide detailed information about synaptic connectivity at the cell compartment level that then rolls up to regional connectivity statements. Second, as neurons are highly polarized cell types, it allows us to assign molecular and functional phenotypes associated with these populations at a granular level.

Finally, because the cell populations in SCKAN are represented using the Neuronal Phenotype Ontology (NPO), they can be compared directly to proposed cell types based on additional phenotypes. The goal of NPO (Gillespie et al., 2022), which forms the backbone of SCKAN, is to represent proposed neuronal types by their phenotypes using a consistent and machine computable representation of molecular, morphological, physiological, anatomical, and phenotypic connectivity characteristics. Using powerful new molecular and correlated techniques, large consortia like the BICCN/BICAN are tackling the open question of defining cell types in the brain (BICCN, 2021). Similar approaches are being employed for the PNS, particularly in the utilization of transcriptomics and spatial transcriptomics to define cell types in peripheral ganglia and cranial nerve nuclei (Bhuiyan et al., 2024). The BICCN cell type mapping of the motor cortex suggests that long range axon projection patterns may not map easily onto transcriptomic and epigenetic cell types, suggesting another level of regulation in defining single-cell connectional specificity (BICCN, 2021). Having a consistent model for modeling phenotypes across these multiple dimensions should facilitate understanding of how proposed cell types map onto connectivity patterns in the PNS.

Despite the potential power of utilizing a neuronal model of connectivity, we did encounter some challenges in utilizing the neuron population as the basis of SCKAN. Defining what constitutes a cell population is somewhat arbitrary: do you represent a population that projects from multiple regions to multiple targets as a single population or as multiple cell populations? In our case, we decided to define a population as a population that subserves a given PNS connection, e.g., the population of pre-ganglionic sympathetic neurons that ultimately enable sympathetic innervation of the heart. So as long as a population could be described as the set of neurons comprising pre-ganglionic sympathetic neurons, the locations of the somata could be distributed. The neuron population approach lends itself to the Flatmap visualization, but creating heat maps and other visualizations of query results is challenging, as querying SCKAN Explorer, for example, returns a list of neuron populations rather than providing a regional connectivity matrix. Thus, the user has to do additional work to find the answer to a simple query such as “What regions project to region X?” As described in a later section, we are developing the next generation of query and visualization tools for SCKAN that will allow users to abstract away the neuron populations to make it easier for them to gain an overview of connectivity for a given query.

### 4.2 Next Steps

The current version of SCKAN supported by the NIH SPARC program was designed as a resource to aid the neuromodulation community through mapping ANS connectivity derived from existing knowledge and new information coming from SPARC. We focused on well established connections in consultation with ANS experts and did not include every experimental study on ANS connectivity or to rare variations or minor projections. Nevertheless, variation across connections, especially at the level of nerve branches is common, as we are seeing in early results from SPARC 2 (Brunsman et al., 2024; Nuzov et al., 2024; Shoffstall et al., 2024). In addition, there are species differences in the origins, routing and termination of various connections (Brauer and Smith, 2015), and inconsistencies across experimental studies of a given connection.

We plan to address these complexities in SCKAN in several ways. At the simplest level, we are planning to use annotation tags to assign a measure of confidence based on the strength of evidence, similar to those used in Hippocampome.org (RRID:SCR 009023). Second, we are developing a quantitative database for SPARC that can be used to compute quantitative information on distributions of cell somas, terminal fields or nerve trajectories for a given neuron population. For example, our current system for assigning location relies on using segmental labels for spinal cord segments and associated DRGs. However, the spinal cord itself is a single column and it is rare to have populations strictly localized. The bulk of a population is often centered on a few segments with the number of neurons dropping off in either direction. Intersegmental routing of axons is also common, where they do not necessarily exist only at the level at which their soma are located. Finally, minor projections are often noted in certain projections, e.g., projections to the ancillary pelvic ganglia from some organs (Keast, 1999).

We are expanding SCKAN to include provenance for individual parts of a neuron population. In most cases, detailed locations of soma, axons and axon terminals are constructed from multiple sources. In some cases we see an inconsistency in a connection between studies for only part of a population. Therefore, we want to annotate individual triples within a given population rather than the entirety of the population. However, current OWL representation requires that annotations be applied to the entire population representation instead of individual parts. To address this, we plan to adopt RDF-star (Arndt et al., 2021), which extends RDF by enabling direct annotation of statements with metadata such as provenance, confidence, or temporal context, eliminating the need for complex reification patterns otherwise required in OWL. SPARQL-star, the query language for RDF-star, streamlines querying annotated data by providing concise and intuitive queries, avoiding verbose OWL-specific patterns, and enhancing usability and efficiency.

Finally, we are developing new interfaces to make it easier for domain experts to explore and visualize the contents of SCKAN. As can be seen from the competency queries presented in the results, creating an all-encompassing interface that is both easy to use yet encapsulates the complexities of SCKAN is not trivial. To date, we have not found a single visualization that allows easy understanding of these basic types of queries. However, we are in the process of replacing the SCKAN Explorer interface, which is designed primarily for curators, with a new tool called SCKANNER. SCKANNER will focus on region-to-region connectivity, abstracting away the individual neuron populations, providing connection matrices, summary reports and the capacity to export search results so that they can be further analyzed. We are also developing new interfaces based on Large Language Models (LLMs) that will allow researchers to query the knowledge base through natural language queries and be provided with human readable summaries of the results. Both of these tools are currently under development but should be released within the next year and will be accessible on the SPARC portal.

### 4.3 Summary and Conclusions

We have described the SCKAN knowledge base, a highly curated knowledge base of detailed connectivity information about the anatomy of the autonomic nervous system. SCKAN is available for query through the SPARC Portal and via a set of programmatic interfaces. SCKAN was designed according to the

FAIR principles, utilizing FAIR vocabularies and building from community standards. As efforts like the BRAIN Initiative CONNECTS program start to generate large amounts of CNS activity, we believe it is important that we design knowledge bases to be interoperable so that they can be combined to provide a comprehensive, machine-readable representation of nervous system connectivity. Moreover, the rapid development of neuromodulation therapies and neuroscience research that extends beyond the brain such as the gut-brain axis suggests that we no longer can relegate the PNS to secondary status.

SCKAN was also designed with an eye towards sustainability, through broad-based support and developing automated and semi-automated tools to help with curation and quality control. All the tools developed by SCKAN are available open source. By utilizing an NLP pipeline and a set of tools for extracting and modeling connectivity information, we ensure that information can be kept up to date, while our neuron-population-based data model makes it possible to include both regional and subcellular connectivity and relate connectivity to transcriptomic and other cell type classifications.

Finally, with generative AI opening up new possibilities for integrating large swaths of human knowledge, resources like SCKAN become important sources of high quality data. Generative AI like LLMs also open up new possibilities for curation, quality control, query and understanding complex knowledge bases. We expect that such capabilities will make knowledge bases such as SCKAN even more important and useful in our quest to understand the nervous system across scales.

## AUTHOR CONTRIBUTIONS

FTI and MEM were the principal authors of the manuscript and key contributors to its conception, design, and coordination. THG and FTI led the overall design and development of SCKAN, including its knowledge engineering, formal modeling, and query mechanisms, and contributed to the development of the supporting tools. THG also made necessary modifications and extensions to NPO for SCKAN, developed and maintained the SCKAN release pipeline, and contributed to manuscript editing.

IZ contributed as the knowledge and data curator, providing essential support to knowledge engineers and assisting with manuscript reviewing and editing. ST contributed to data collection, manuscript drafting and editing, and participated in the development of the Composer and SCKANNER interfaces. MS-Z managed the InterLex term request pipeline for SCKAN, served as the primary curator for the translation of the ApiNATOMY models, and contributed to manuscript reviewing and editing.

BO designed and developed the NLP pipeline for SCKAN and provided related statistics. BdB, as the primary author of the ApiNATOMY models, contributed to the initial SCKAN models and provided feedback on the manuscript. JB, as the overall project manager for SCKAN, significantly contributed to manuscript review and editing. JSG contributed to the design and development of curatorial tools and provided feedback. MEM is the principal investigator of the project. All authors read and approved the final manuscript.

## ACKNOWLEDGMENTS

The authors thank members of the SPARC MAP-CORE, including David Brooks, Yuda Munarko, and David Nickerson, for helpful discussions and SCKAN model testing. The SPARC MAP-CORE is led by Peter Hunter, FRS, FRSNZ, and David Nickerson, PhD, at the Auckland Bioengineering Institute/Pūhanga Koiora o Tāmaki Makaurau), University of Auckland/Waipapa Taumata Rau.

## DATA AND CODE AVAILABILITY STATEMENT

*•* Latest SCKAN Release: https://doi.org/10.5281/zenodo.5337441
*•* Accessing SCKAN: Link to the documentation
*•* About Simple SCKAN: https://github.com/SciCrunch/sparc-curation/tree/master/docs/simple-sckan
*•* SPARQL code and results used for the CQs in this paper: Link to the repository
*•* Example Queries in Jupyter Notebook: https://github.com/smtifahim/sckan-query-examples
*•* Published ApINATOMY models: https://doi.org/10.5281/zenodo.5519557

